# The *Magnaporthe oryzae* MAX effector AVR-Pia binds a novel group of rice HMA domain-containing proteins

**DOI:** 10.1101/2025.07.11.664054

**Authors:** Josephine H.R. Maidment, Svenja C. Saile, Aurélien Bocquet, Céline Thivolle, Léo Bourcet, Lisa-Fatimatou Planel, Muriel Gelin, Thomas Kroj, André Padilla, Karine de Guillen, Stella Cesari

## Abstract

Phytopathogenic fungi secrete effector proteins to promote virulence. MAX (*<u>M</u>agnaporthe* <u>A</u>vrs and <u>T</u>oxB-like) effectors form a sequence-diverse family sharing a conserved protein structure. AVR-Pia, a MAX effector from the rice blast fungus *Magnaporthe oryzae,* is recognised by the paired rice nucleotide-binding leucine-rich repeat immune receptors OsRGA4/OsRGA5 through direct binding to an integrated heavy metal-associated (HMA) domain in OsRGA5. Here, we identify previously unknown host targets of AVR-Pia: four HMA domain-containing rice proteins, belonging to the <u>H</u>MA Isoprenylated <u>P</u>lant <u>P</u>roteins (HIPPs) and <u>H</u>MA <u>P</u>lant <u>P</u>roteins (HPPs). AVR-Pia interacted with all four proteins, both in vitro and in planta, and bound their HMA domains with varying affinities. The crystal structure of AVR-Pia in complex with the HMA domain of OsHPP09 revealed the molecular details of the binding interface. Structure-guided mutagenesis of OsHPP09 identified a single point mutation which prevents AVR-Pia binding, providing a foundation for targeted engineering of HMA domains to evade effector binding.

## Introduction

The fungal pathogen *Magnaporthe oryzae* (syn. *Pyricularia oryzae*) is the causative agent of rice blast, one of the most devastating diseases affecting cultivated rice worldwide^1,2^. *M. oryzae* also causes severe yield losses in other cereals, including millets, barley and wheat, threatening global food security^1^.

To successfully colonise host plants, *M. oryzae* must evade plant defence responses and manipulate host cellular pathways to its advantage. Central to infection is the secretion of a large repertoire of effector proteins, many of which can be categorised into structural families^3^. In *M. oryzae,* the *<u>M</u>agnaporthe* <u>A</u>VRs and To<u>x</u>B-like (MAX) effectors form the largest structural family^4^. MAX effectors are defined by a conserved fold consisting of two β-sheets, each composed of three antiparallel β-strands and stabilised by one or more disulphide bonds. Despite this shared structure, there is little amino acid sequence similarity between MAX effectors. They differ in surface properties and some carry variable N- and C-terminal extensions, which can influence their folding pathway^5^ and mediate interactions with host proteins^3,6–9^.

AVR-Pia is a MAX effector present in approximately 20% of rice-infecting *M. oryzae* isolates. It is highly expressed during the biotrophic phase of host colonisation and contributes to virulence on rice^4,7,10^. AVR-Pia can be recognised by the intracellular immune receptor OsRGA5, a nucleotide-binding leucine-rich repeat (NLR) protein that functions together with the helper NLR OsRGA4 to activate immune responses and restrict the spread of *M. oryzae*^11–13^. Canonical plant NLRs consist of an N-terminal signalling domain, a central nucleotide-binding domain and a C-terminal leucine-rich repeat domain^14^. However, many NLRs also carry non-canonical integrated domains (IDs), thought to originate from host proteins targeted by pathogen effectors^15^. These IDs, found across diverse plant species, are believed to function as mimics or baits for effector recognition (“integrated decoy” model)^15–18^. Despite sharing limited sequence similarity, both AVR-Pia and a second MAX effector, AVR1-CO39, bind to the heavy metal-associated (HMA) ID of OsRGA5^11,12^. While this suggests that these effectors may target host proteins containing HMA domains to promote disease, no host targets of AVR-Pia or AVR1-CO39 have been identified, and their specific virulence functions remain unknown.

The OsRGA5 HMA ID shares sequence similarity with a family of HMA domain-containing metallochaperone-like proteins. The family can be subdivided into two groups: <u>H</u>MA domain-containing Isoprenylated <u>P</u>lant <u>P</u>roteins (HIPPs) that carry a C-terminal CaaX isoprenylation motif, and <u>H</u>MA domain-containing <u>P</u>lant <u>P</u>roteins (HPPs) that lack this motif^19,20^. The two groups are collectively referred to hereafter as H(I)PPs. Isoprenylation, a post-translational modification involving the covalent attachment of an isoprenoid lipid to the cysteine residue in the CaaX motif, has been reported to facilitate membrane association^21^. Many HIPPs harbour proline-rich regions between the HMA domain and the CaaX motif, potentially mediating protein-protein interactions^19,22^. Typically, HMA domains feature a metal-binding MxCxxC motif, implicating H(I)PPs in roles related to metal homeostasis^23,24^. However, approximately a third of rice H(I)PPs lack one or both cysteines in their MxCxxC motif, suggesting functional diversification beyond metal binding. H(I)PPs have undergone extensive expansion in vascular plants, with approximately 100 members identified in rice^19,25–27^. Most remain functionally uncharacterized, although several have been implicated in response to various biotic and abiotic stresses^19,28–30^. Increasing evidence points to a broader role of H(I)PPs as susceptibility factors in interactions with pathogens, pests, and viruses^27,31–41^.

In cereals, H(I)PPs have been identified as susceptibility factors for *M. oryzae* infection^27,33,34^. A loss-of-function allele of *OsHIPP05* (*Pi21*) contributes to blast resistance in the field. *OsHIPP20* is also a susceptibility gene; a loss-of-function mutation reduces the susceptibility of rice to *M. oryzae*^27^. Some H(I)PPs are targeted by *M. oryzae* MAX effectors AVR-Pik and APikL2^8,27,42^. AVR-Pik specifically interacts with members of a phylogenetic subclade of OsH(I)PPs, referred to as clade A, including OsHIPP19, OsHIPP20, OsHPP03, and OsHPP04^8,27^. The paired rice NLR proteins OsPik-1 and OsPik-2 mediate recognition of AVR-Pik, which requires direct binding of the effector to an HMA ID in OsPik-1^43–45^. The HMA domain of OsPik-1 belongs to the same phylogenetic clade as the H(I)PPs targeted by AVR-Pik, reinforcing the idea that effector targets may serve as templates for domain integration into NLRs.

Structural analyses of complexes between AVR-PikF or APikL2 and the HMA domains of their HIPP targets or cognate NLR receptors have revealed a conserved binding interface centring on β3 of the MAX fold and β4 of the HMA domain^8,42,45,46^. By contrast, the crystal structure of AVR1-CO39 bound to the HMA ID of OsRGA5 revealed a markedly different interface involving β2 of both AVR1-CO39 and OsRGA5-HMA^12^. NMR titration experiments, structural modelling, and mutagenesis assays indicated that AVR-Pia binds OsRGA5-HMA at a similar interface^12,13,47^. Additionally, the crystal structure of AVR-Pia in complex with Os-Pikp-HMA revealed a similar binding interface, indicating that AVR-Pia binds different HMA domains in a structurally similar manner^48^. The crystal structure of the MAX effector Pwl2 bound to OsHIPP43 also revealed an interface involving β2 of the Pwl2 MAX fold and β2 of OsHIPP43-HMA^9^. However, unlike AVR-Pia and AVR1-CO39, Pwl2 has a C-terminal extension, comprising an ⍺-helix and an unstructured region, which contributes significantly to HMA binding^9^. Collectively, these structural studies underscore the remarkable diversity with which MAX effectors have evolved to bind various HMA domain-containing proteins. They support the hypothesis that AVR-Pia and AVR1-CO39 target distinct H(I)PPs through interfaces that differ from those employed by other MAX effectors.

Here, we identified four OsH(I)PPs as specific interactors of AVR-Pia, none of which are bound by the MAX effectors AVR1-CO39, AVR-Pik and Pwl2. Using biochemical and biophysical approaches, we characterised the interaction between AVR-Pia and one of the interactors, OsHPP09. The crystal structure of AVR-Pia/OsHPP09-HMA complex showed global similarity to the AVR1-CO39/OsRGA5-HMA complex but revealed more extensive intermolecular hydrogen bonding. Guided by the AVR-Pia/OsHPP09-HMA structure, we introduced a point mutation in OsHPP09 which disrupts its interaction with AVR-Pia without compromising the HMA fold.

## Results

### AVR-Pia interacts with rice H(I)PPs in Y2H screening

As AVR-Pia is recognised by OsRGA5 through direct binding to its HMA ID^11,13^, we hypothesised that AVR-Pia may target HMA domain-containing proteins to promote host susceptibility. We conducted pairwise yeast two-hybrid (Y2H) assays, using a signal peptide-deleted version of AVR-Pia (dSP-AVR-Pia, 20-85 aa) fused to the GAL4 activation domain (AD) against a rice HMA domain library^49^ comprising 78 rice H(I)PP HMA domains fused to the GAL4 DNA binding domain (BD). Five HMA domains interacted with AVR-Pia: OsHPP09 (LOC_Os03g02070), OsHIPP41 (LOC_Os03g06080), OsHPP11 (LOC_Os04g45130), OsHIPP14 (LOC_Os04g39320) and OsHIPP39 (LOC_Os03g02860) (**Fig. 1a, Supplementary Fig. 1, Supplementary Table 1**). When paired with AVR-Pia, OsHPP09-HMA conferred strong yeast growth on interaction-selective media, indicating a robust interaction, while yeast growth for the remaining candidates was weaker.

**Fig. 1.**
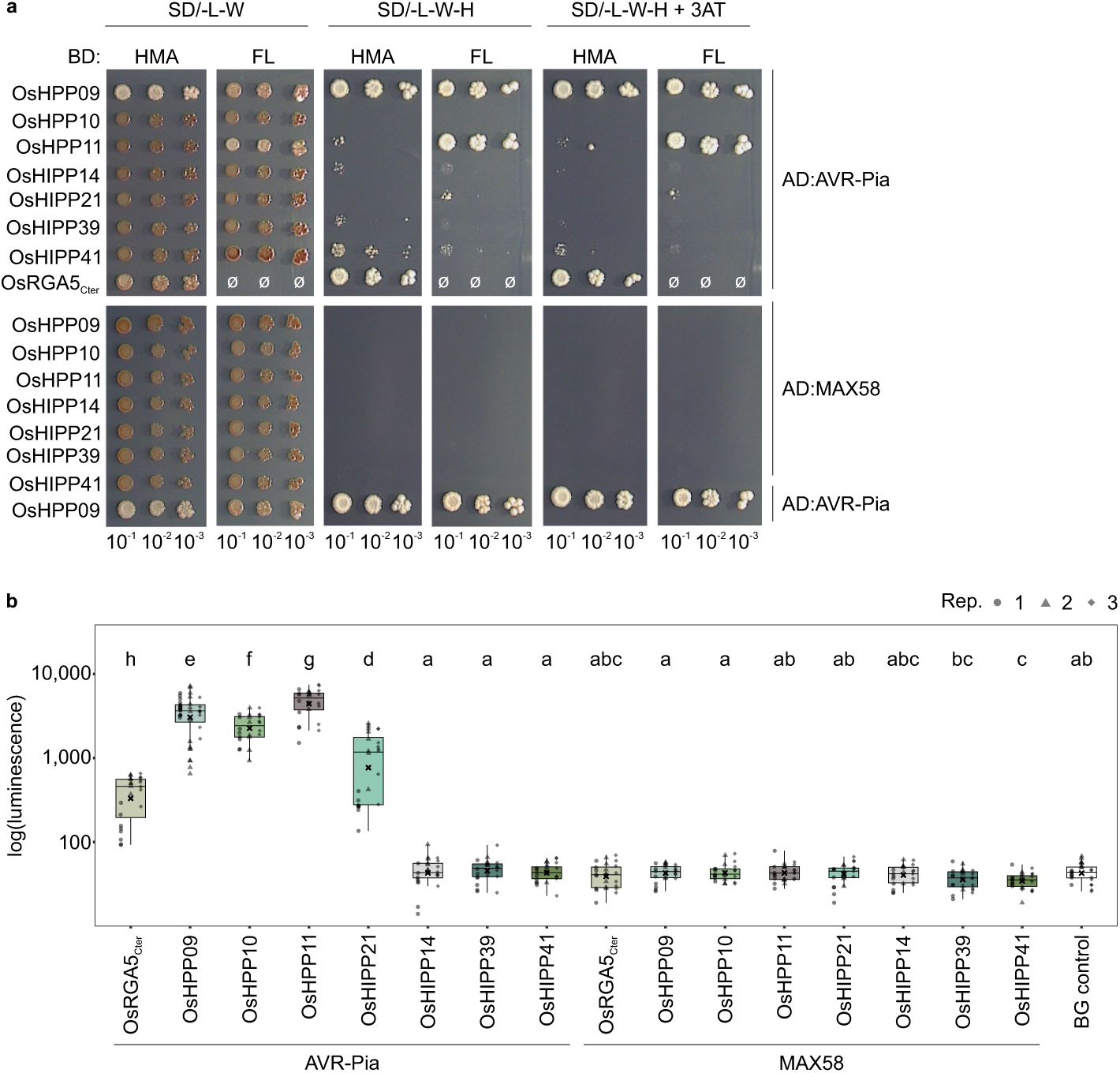
AVR-Pia interacts with OsH(I)PPs in yeast two-hybrid and in planta. **A** Y2H assay of AVR-Pia and MAX58 (both without signal peptides, ΔSP) with either the HMA domain of OsH(I)PPs or full-length (FL) proteins. AVR-Pia/OsRGA5_Cter_ (883-1116 aa) and AVR-Pia/OsHPP09 served as positive controls (upper and lower panels, respectively). Serial dilutions of diploid yeast clones were spotted onto synthetic defined (SD) media to monitor growth (SD/-LW), or to assess protein-protein interactions (SD/-LWH and SD/-LWH supplemented with 0.5 mM 3-amino-1,2,4-triazole (3AT)). Photos were taken after 7 days of incubation. Ø indicates positions where no yeast was spotted. AD, GAL4 activation domain; BD, GAL4 DNA binding domain. **b** Split luciferase complementation assay of AVR-Pia and MAX58 (both ΔSP; fused to the N-terminal part of luciferase; 3xHA:NLuc:MAX) with FL OsH(I)PPs (fused to the C-terminal part of luciferase: 3xFlag:CLuc:OsH(I)PP). Indicated constructs were transiently co-expressed in *N. benthamiana* leaves and leaf discs were harvested 2 days post-infiltration for luminescence measurements. AVR-Pia/OsRGA5_Cter_ (883-1116 aa) was used as a positive control, and leaves infiltrated with P19 alone as background (BG) control. Box plots show the median (line), mean (cross) and upper/lower quartiles (box limits), whiskers extend to the most extreme data points within 1.5x the interquartile range, with outliers plotted individually. Three independent replicates were performed (n = 8 per replicate) except for AVR-Pia/OsHPP09: n = 16 in replicates 1 and 2. Different letters indicate statistically significant differences based on a pairwise Wilcoxon test (α = 0.05).

We additionally tested the HMA library against the well-characterised MAX effectors Pwl2 and AVR-PikD. Consistent with earlier publications, we observed dSP-Pwl2 (22-145 aa) interacting with the HMA domain of OsHIPP43, and dSP-AVR-PikD (22-113 aa) interacting with the HMA domains of OsHIPP19, OsHIPP20, and other phylogenetically related HMA domains^8,9,27,34^ (**Supplementary Fig. 1**), confirming the sensitivity of our screen.

In parallel, we performed an independent Y2H screen using a rice cDNA library. This identified OsHIPP21 (LOC_Os09g09830) as a putative interactor of AVR-Pia (**Supplementary Table 1, Supplementary Table 2**).

For each interacting HMA domain, we next tested whether full-length (FL) proteins also bind AVR-Pia in pairwise Y2H assays. Interestingly, interaction between AVR-Pia and FL OsH(I)PP proteins was only observed for OsHPP09, OsHPP11 and OsHIPP21 (**Fig. 1a and Supplementary Fig. 2**), despite confirmed expression of all proteins (**Supplementary Fig. 3a, 3b, 3c**). To exclude the possibility that the isoprenylation (CaaX) motif of studied FL HIPPs affects nuclear localisation in yeast^21^, we mutated the cysteine to serine (**Supplementary Fig. 3a, 3b**). However, OsHIPP14^C187S^, OsHIPP39^C190S^ and OsHIPP41^C152S^ did not interact with AVR-Pia in Y2H assays (**Supplementary Fig. 2**).

The HMA domains of OsHPP09 and OsHPP11 share 91.5 % and 74.6 % amino acid sequence identity with OsHPP10 (LOC_Os10g36200) (**Supplementary Fig. 4**). Surprisingly, AVR-Pia did not interact with OsHPP10-HMA nor the FL protein in Y2H assays (**Fig. 1a**). By contrast, OsHIPP21-HMA shares only 30-40% sequence identity with the HMA domains of the three OsHPPs (**Supplementary Fig. 4**), and while the HMA domain of OsHIPP21 did not interact with AVR-Pia, FL OsHIPP21 did. Taken together, these results suggest that AVR-Pia binds and potentially targets OsHPP09, OsHPP11, and OsHIPP21.

Previous work classified MAX effectors into 20 structural groups^6^. MAX58 belongs to the same group as AVR-Pia, but differs in surface properties^6^. dSP-MAX58 (20-86 aa) did not interact with any of the tested OsH(I)PPs in Y2H assays (**Fig. 1a, Supplementary Fig. 1**), although the effector protein was expressed (**Supplementary Fig. 3c**), suggesting that, despite their shared structural fold, effectors with distinct surface properties target different host proteins.

### AVR-Pia interacts with rice H(I)PPs in planta

To test whether AVR-Pia and the four OsH(I)PPs interact in planta, we performed split-luciferase complementation assays in *Nicotiana benthamiana*. 3xFlag-CLuc-OsH(I)PP FL proteins were co-expressed with 3xHA-NLuc-dSP-AVR-Pia or 3xHA-NLuc-dSP-MAX58. Co-expression of FL OsHPP09, OsHPP11, and OsHIPP21 with AVR-Pia led to high luminescence levels, exceeding those observed for the positive control OsRGA5_Cter_ (883-1116 aa, including the HMA ID) (**Fig. 1b**). Strikingly, a strong luminescence signal was observed for OsHPP10/AVR-Pia, within a similar range to OsHPP09 and OsHPP11 and exceeding OsHIPP21 (**Fig. 1b**). Consistent with the Y2H results, no interaction was observed in planta between AVR-Pia and other tested FL OsHIPPs, including OsHIPP14, OsHIPP39 and OsHIPP41 (**Fig. 1b**). MAX58 showed no interaction with any of the tested OsH(I)PPs (**Fig. 1b**). Expression of all proteins was confirmed by immunoblot analyses (**Supplementary Fig. 5**). These data demonstrate that AVR-Pia interacts in planta with OsHPP09, OsHPP10, OsHPP11 and OsHIPP21.

### AVR-Pia interacts with the HMA domains of OsHPP09, OsHPP10, OsHPP11 and OsHIPP21 in vitro with varying affinities

Recombinant AVR-Pia and HMA domains were produced and purified from *E. coli* (**Supplementary Fig. 6**). We first tested for interaction between purified OsHPP09-HMA and AVR-Pia using analytical size exclusion chromatography. When analysed alone, the peak elution volumes for AVR-Pia and OsHPP09-HMA were 15.3 ml and 13.0 ml, respectively (the latter is consistent with dimerisation, as reported for other HMA domains in solution^12,45^). Following incubation with OsHPP09-HMA (1:1 effector:HMA ratio), the peak elution volume of AVR-Pia shifted to 12.7 ml, consistent with formation of an AVR-Pia/OsHPP09-HMA complex (**Supplementary Fig. 7**). The co-elution of the two proteins was supported by SDS-PAGE analysis of elution fractions (**Supplementary Fig. 7**).

To determine the affinity with which AVR-Pia binds to the HMA domains of OsHPP09, OsHPP10, OsHPP11 and OsHIPP21, we used isothermal titration calorimetry (ITC). Titration of AVR-Pia into a solution containing any of the purified HMA domains resulted in negative peaks in the titration curve (**Fig. 2, Supplementary Fig. 8**), indicating an exothermic binding interaction. Dissociation constant (*K*_D_) values were determined from binding isotherms (**Fig. 2, Supplementary Fig. 8**) fitted to a single site model.

**Fig. 2.**
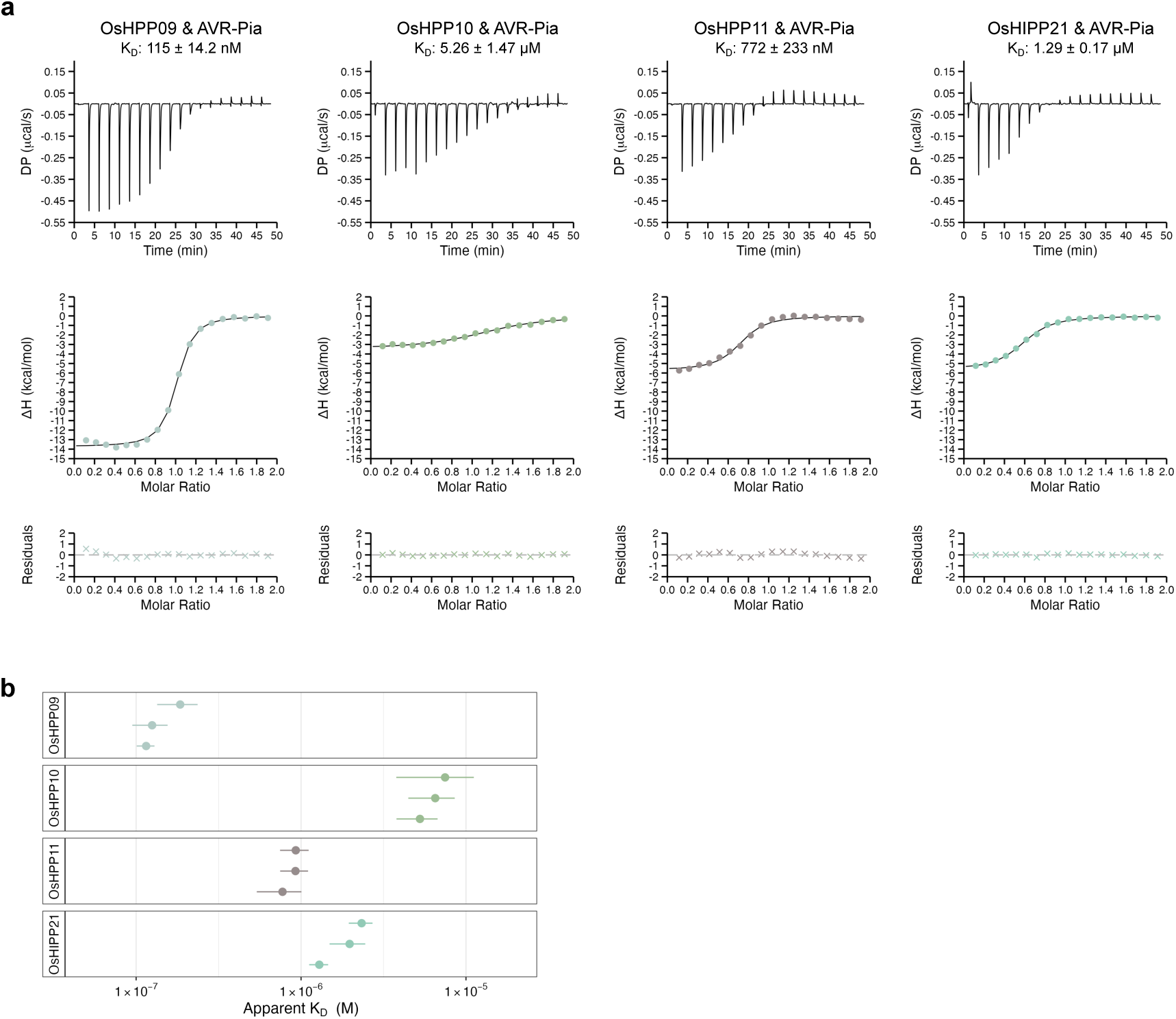
AVR-Pia binds the HMA domains of OsH(I)PPs with varying affinities. **a** Representative ITC experiments for each of the AVR-Pia/HMA interactions tested. Top panel shows the raw thermogram obtained from titration of AVR-Pia into a solution containing the purified HMA domains. Central panel shows the integrated heats (coloured dots) and binding isotherm fitted to a single site model (black line) using the MicroCal PEAQ-ITC analysis software (Malvern Panalytical). Bottom panel shows the differences (coloured crosses) between the modelled and observed values (residuals). **b** Dot plot representation of the apparent *K*_D_ for each of the AVR-Pia/HMA interactions in three independent experiments. Coloured dots represent the *K*_D_ values determined in each replicate with coloured lines representing the error associated with the *K*_D_ value as determined by the MicroCal PEAQ-ITC analysis software (Malvern Panalytical).

AVR-Pia bound to OsHPP09-HMA with an apparent *K*_D_ of 115 nM (**Fig. 2**). Despite differing in just six amino acids, AVR-Pia bound OsHPP10-HMA with significantly lower affinity, with an apparent *K*_D_ of 5.26 µM. The binding affinities of AVR-Pia and OsHPP11-HMA or OsHIPP21-HMA fell between these two extremes, with apparent *K*_D_s of 772 nM and 1.29 µM, respectively (**Fig. 2, Supplementary Fig. 8, Supplementary Fig. 9, Supplementary Table 3**). The apparent *K*_D_s, determined by ITC, for the interactions between OsRGA5-HMA and AVR-Pia or AVR1-CO39 were previously reported as 7.8 µM^13^ and 5.1 µM^12^, respectively.

### AlphaFold Multimer models cannot differentiate between HMA domains which are bound by AVR-Pia and those which are not

Several studies have used AlphaFold Multimer to distinguish interacting from non-interacting pairs of effectors/host proteins^27,50,51^. We therefore tested whether structural models of AVR-Pia in complex with experimentally verified interactors (OsHPP09-HMA, OsHPP10-HMA, OsHPP11-HMA and OsHIPP21-HMA) showed differences, either in the models themselves or in the associated confidence metrics, compared to structural models of AVR-Pia with non-interacting HMA domains. We modelled AVR-Pia/HMA complexes using AlphaFold2 and AlphaFold3. High confidence models were obtained for all AVR-Pia/HMA pairs except AVR-Pia/OsHIPP42-HMA using AlphaFold2 (**Supplementary Fig. 10**) and AVR-Pia/OsHIPP41-HMA using AlphaFold3 (**Supplementary Fig. 11**). For the AlphaFold2 models, interface analysis with qtPISA revealed a tendency towards increased interface area for AVR-Pia-interacting HMA domains compared to non-interacting HMA domains. (**Supplementary Fig. 10)**. By contrast, there were no significant differences for AlphaFold3 models (**Supplementary Fig. 11**). Overall, AlphaFold could not clearly differentiate between HMA domains bound or not bound by AVR-Pia.

### AVR-Pia-H3 does not interact with OsHPP09, OsHPP10, OsHPP11 and OsHIPP21

Some rice-infecting *M. oryzae* isolates carry an allele of AVR-Pia, AVR-Pia-H3, which contains two non-synonymous polymorphisms (F24S and T46N)^11^ (**Supplementary Fig. 12**). These polymorphisms interfere with OsRGA5-HMA binding and allow evasion of OsRGA4/OsRGA5-mediated immunity^11,13^. Pairwise Y2H assays revealed that AVR-Pia-H3 failed to interact with FL OsHPP09, OsHPP10, OsHPP11 and OsHIPP21 or their HMA domains (**Fig. 3a**). In planta split-luciferase complementation assays gave only weak luminescence signals for AVR-Pia-H3 with OsH(I)PPs, comparable to the negative control pairs OsRGA5_Cter_/AVR-Pia-H3 and OsHPP09/MAX58, and markedly lower than the positive control OsHPP09/AVR-Pia (**Fig. 3b**). Protein expression in yeast and in planta was confirmed by immunoblot (**Supplementary Fig. 3c and 13**). Analysis of a 1:1 effector:HMA mixture of purified recombinant AVR-Pia-H3 and OsHPP09-HMA by analytical size exclusion chromatography showed no co-elution of the two proteins, indicating that they did not form a stable complex in vitro (**Fig. 3c**).

**Fig. 3.**
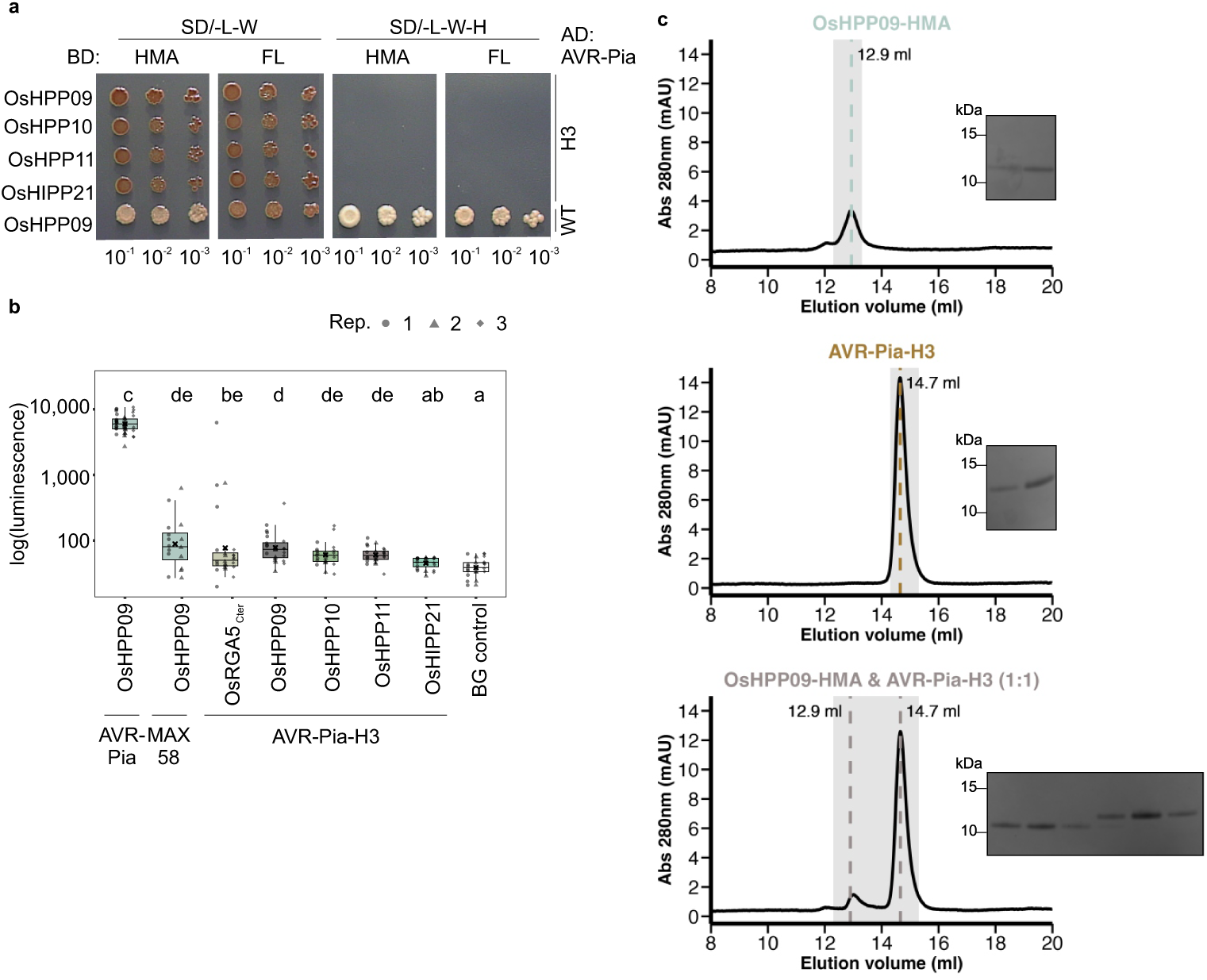
AVR-Pia-H3 does not interact with OsHPP09, OsHPP10, OsHPP11 and HIPP21. **a** Y2H interaction analysis between AVR-Pia-H3 (ΔSP) and either the HMA domain of OsH(I)PPs or full-length (FL) proteins. OsHPP09/AVR-Pia served as positive control. Serial dilutions of diploid yeast were spotted onto synthetic defined (SD) media to monitor growth (SD/-LW) or to assess protein-protein interactions (SD/-LWH). Photos were taken after 7 days of incubation. AD, GAL4 activation domain; BD, GAL4 DNA binding domain. **b** Split luciferase complementation assay of AVR-Pia-H3 (ΔSP; fused to the N-terminal part of luciferase; 3xHA:NLuc:MAX) with FL OsH(I)PPs (fused to the C-terminal part of luciferase: 3xFlag:CLuc:OsH(I)PP). Indicated constructs were transiently co-expressed in *N. benthamiana* leaves and leaf discs were harvested 2 days post-infiltration for luminescence measurements. AVR-Pia/OsHPP09 served as a positive control, MAX58/OsHPP09 and AVR-Pia-H3/OsRGA5_Cter_ (883-1116 aa) as negative controls, and leaves infiltrated with P19 alone as background (BG) control. Box plots show the median (line), mean (cross) and upper/lower quartiles (box limits), whiskers extend to the most extreme data points within 1.5x the interquartile range, with outliers plotted individually. Three independent replicates were performed, with n = 8 per combination per replicate. Due to experimental constraints, for AVR-Pia/OsHPP09, n = 16 in replicates 1 and 2. Data points are shown with shapes indicating the replicate. Different letters indicate statistically significant differences based on a pairwise Wilcoxon test (α = 0.05). **c** Analytical gel filtration traces obtained from injection of OsHPP09-HMA alone (top panel), AVR-Pia-H3 alone (middle panel) and the proteins combined (1:1 molar ratio; bottom panel). Significant peaks are indicated by coloured dashed lines with elution volume labelled. SDS-PAGE gel inserts show fractions from peak elution volumes indicated by grey shaded regions. OsHPP09-HMA absorbs light at 280 nm poorly (molar extinction coefficient: 1490 M^-1^cm^-1^) so the peak corresponding to OsHPP09-HMA is small.

Together, these data indicate that the F24S and T46N polymorphisms that disrupt binding of AVR-Pia to the HMA ID of OsRGA5 also interfere with binding to its candidate host targets.

### AVR1-CO39 does not interact with OsHPP09, OsHPP10, OsHPP11 and OsHIPP21

As the HMA ID of OsRGA5 also binds the MAX effector AVR1-CO39^11,12^, we tested whether AVR1-CO39 binds to the same H(I)PPs as AVR-Pia. dSP-AVR1-CO39 (22-89 aa) did not interact with FL OsHPP09, OsHPP10, OsHPP11 or OsHIPP21, nor their HMA domains, in Y2H, whereas interaction with OsRGA5_Cter_ was confirmed (**Fig. 4a**). Effector expression was confirmed by immunoblot (**Supplementary Fig. 3c**).

**Fig. 4.**
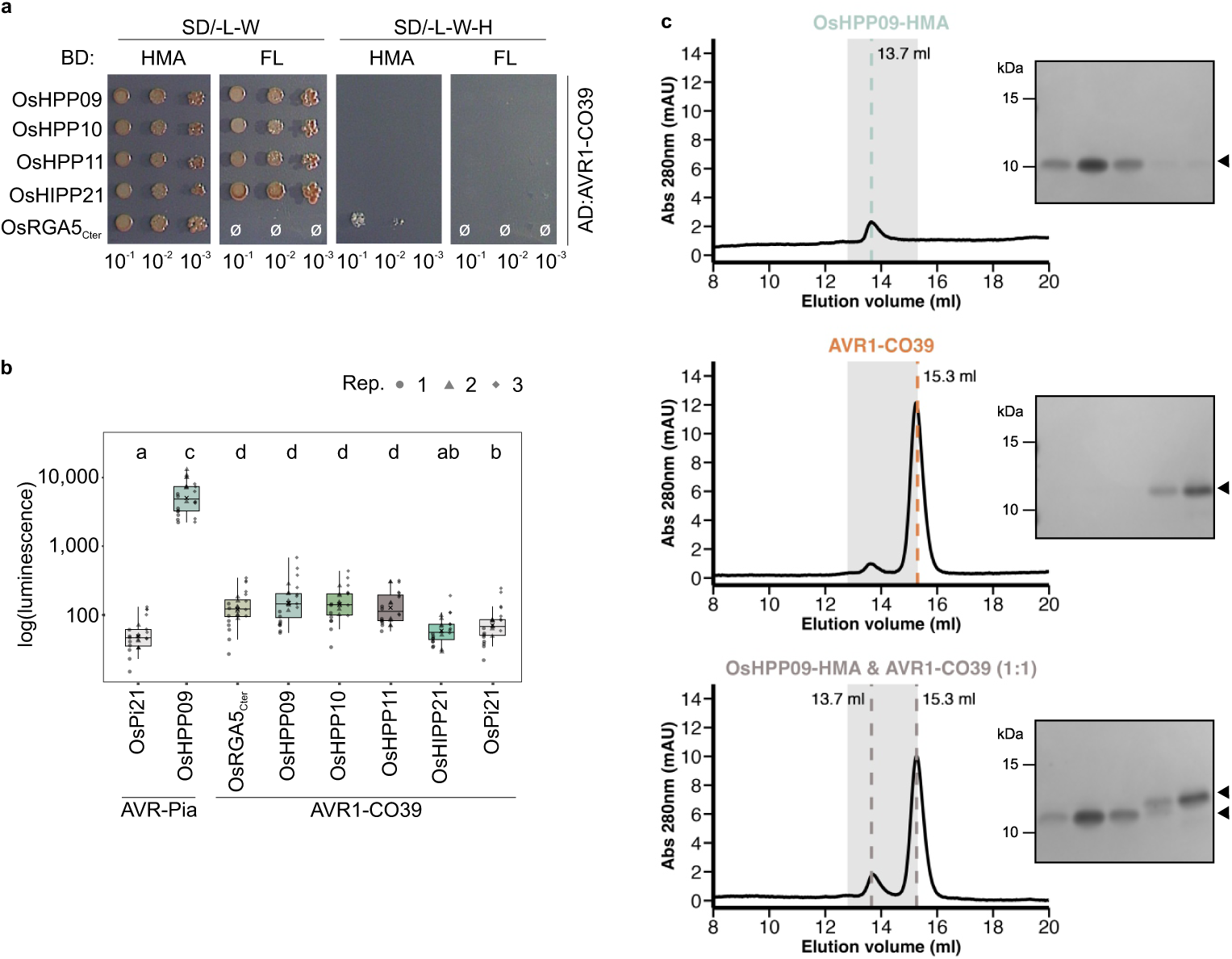
OsH(I)PPs interacting with AVR-Pia do not interact with AVR1-CO39. **a** Y2H interaction analysis between AVR1-CO39 (ΔSP) and either the HMA domain of OsH(I)PPs or full-length (FL) proteins. AVR1-CO39/OsRGA5_Cter_ (883-1116 aa) served as a positive control. Serial dilutions of diploid yeast were spotted onto synthetic defined (SD) media to monitor growth (SD/-LW) or to assess protein-protein interactions (SD/-LWH). Photos were taken after 7 days of incubation. Ø indicates positions where no yeast was spotted. AD, GAL4 activation domain; BD, GAL4 DNA binding domain. **b** Split-luciferase complementation assay of ΔSP AVR1-CO39 with FL OsH(I)PPs and OsRGA5_Cter_ (883-1116 aa). MAX effectors were fused to the N-terminal parts of luciferase (3xHA:NLuc:MAX), while OsH(I)PP FL proteins and OsRGA5_Cter_ were fused to its C-terminal part (3xFlag:CLuc:OsH(I)PP/OsRGA5_Cter_). Constructs were transiently co-expressed in *N. benthamiana* leaves and leaf discs were harvested 2 days post-infiltration for luminescence measurements. AVR-Pia or AVR1-CO39 with OsPi21 served as negative controls and AVR-Pia/OsHPP09 as a positive control. Box plots show the median (line), mean (cross) and upper/lower quartiles (box limits), whiskers extend to the most extreme data points within 1.5x the interquartile range, with outliers plotted individually. Three independent replicates were performed, with n = 8 per combination per replicate. All data points are shown with shapes indicating the replicate. Different letters indicate statistically significant differences based on a pairwise Wilcoxon test (α = 0.05). **c** Analytical gel filtration traces obtained from injection of OsHPP09-HMA alone (top panel), AVR1-CO39 alone (middle panel) and the proteins combined (1:1 molar ratio; bottom panel). Significant peaks are indicated by coloured dashed lines with elution volume labelled. SDS-PAGE gel inserts show fractions from peak elution volumes indicated by grey shaded regions. The small peak at ∼13.7 ml for AVR1-CO39 is likely due to dimerisation of the AVR1-CO39 protein; the quantity of the dimer is insufficient to observe a band on the SDS-PAGE gel. OsHPP09-HMA absorbs light at 280 nm poorly (molar extinction coefficient: 1490 M^-1^cm^-1^) so the peak corresponding to OsHPP09-HMA is small.

In split-luciferase complementation assays, we observed weak luminescence for AVR1-CO39 with FL OsHPP09, OsHPP10 and OsHPP11, similar to the luminescence observed with OsRGA5_Cter_ but substantially lower than the level observed for AVR-Pia/OsHPP09 (**Fig. 4b**). Consistent with the Y2H results, no interaction was detected between FL OsHIPP21 and AVR1-CO39, similar to the negative control OsPi21/AVR1-CO39 (**Fig. 4b**). Protein expression was validated by immunoblot (**Supplementary Fig. 14**). Analytical size-exclusion chromatography with purified proteins further supported the absence of complex formation (**Fig. 4c**).

Collectively, these results suggest that, while AVR1-CO39 and AVR-Pia are both recognised by the same HMA ID in OsRGA5, they do not bind the same H(I)PPs.

### AVR-Pia interacts with a distinct group of H(I)PPs to AVR-Pik and Pwl2

Previous studies have shown that the MAX effectors AVR-PikC, AVR-PikD and Pwl2 target H(I)PPs^8,9,27,34^. However, dSP-AVR-PikC (22-113 aa), dSP-AVR-PikD (22-113 aa) and dSP-Pwl2 (22-145 aa) did not bind to the AVR-Pia interactors OsHPP09, OsHPP10, OsHPP11 and OsHIPP21 in Y2H assays (**Fig. 5a**), while we confirmed the known interactions of AVR-PikC/AVR-PikD with OsHIPP19/OsHIPP20, and Pwl2 with OsHIPP43 (**Fig. 5a**). Interestingly, AVR-Pia interacted with FL OsHIPP19 under low-stringency Y2H conditions (**Fig. 5a**).

**Fig. 5.**
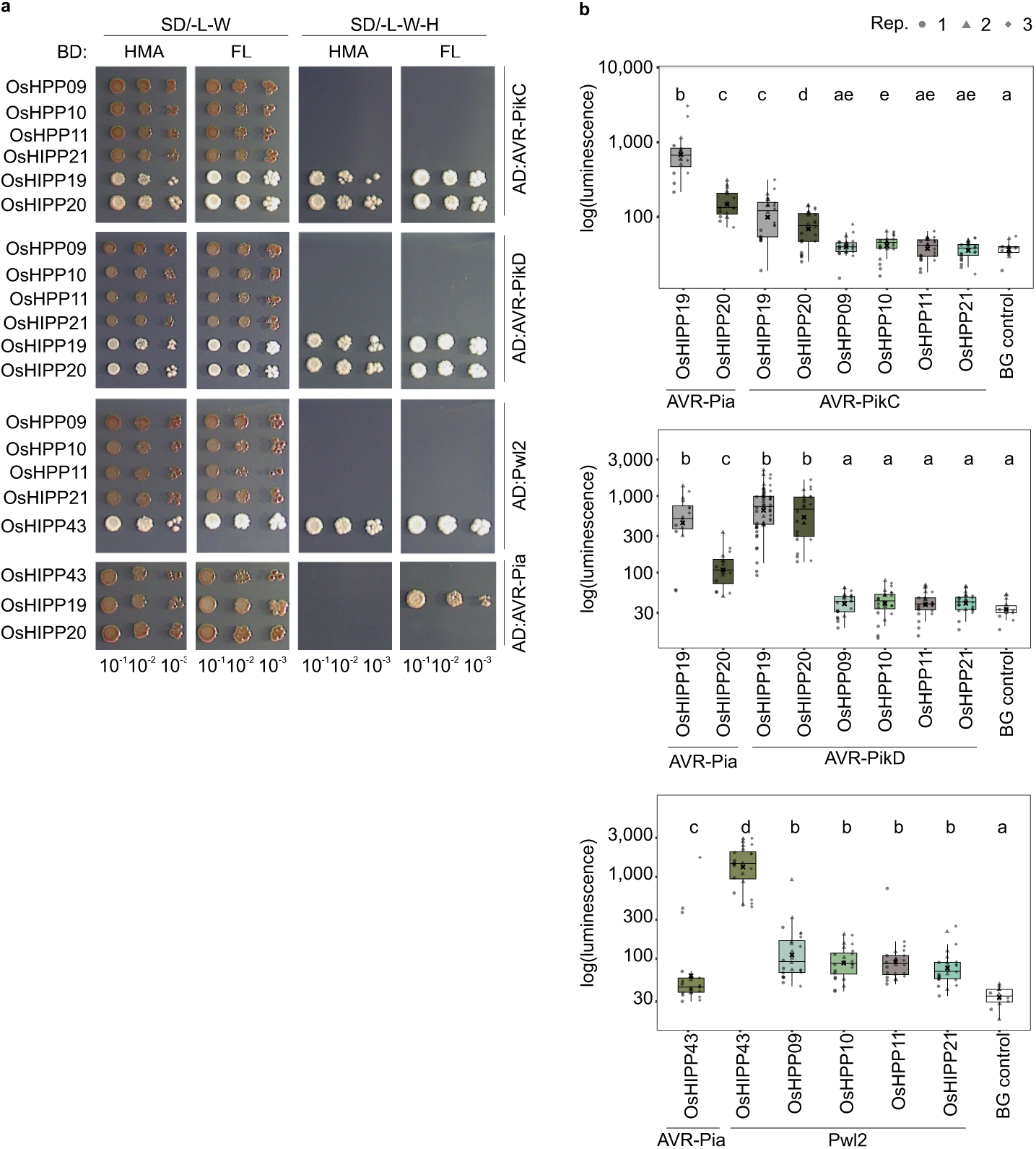
OsH(I)PPs interacting with AVR-Pia do not interact with AVR-Pik or Pwl2. **a** Y2H interaction analysis between AVR-PikC, AVR-PikD, Pwl2 and AVR-Pia (all ΔSP) and either the HMA domain of OsH(I)PPs or full-length (FL) proteins. OsHIPP19/AVR-PikC, OsHIPP20/AVR-PikC, OsHIPP19/AVR-PikD, OsHIPP20/AVR-PikD and OsHIPP43/Pwl2 served as positive controls. Serial dilutions of diploid yeast were spotted onto synthetic defined (SD) media to monitor growth (SD/-LW) or to assess protein-protein interactions (SD/-LWH). Photos were taken after 7 days of incubation. AD, activating domain; BD, binding domain. **b** Split luciferase complementation assay of MAX effectors (ΔSP; fused to the N-terminal part of luciferase: 3xHA:NLuc:MAX) with OsH(I)PP FL proteins (fused to the C-terminal part of luciferase: 3xFlag:CLuc:OsH(I)PP). AVR-PikC/AVR-PikD with OsHIPP19/20 and Pwl2 with OsHIPP43 served as positive controls, AVR-Pia with OsHIPP43 served as a negative control. Leaves infiltrated with P19 alone served as background (BG) control. Box plots show the median (line), mean (cross) and upper/lower quartiles (box limits), whiskers extend to the most extreme data points within 1.5x the interquartile range, with outliers plotted individually Three independent replicates were performed, with n = 8 per combination per replicate, except for AVR-Pia/OsHIPP19 and AVR-Pia/OsHIPP20 in AVR-PikC and AVR-PikD panels: n = 4 in replicates 1 and 3. For BG control in AVR-PikC and AVR-PikD panels: n = 3 in replicates 1 and 3 and for BG control in Pwl2 panel: n = 2 in replicates 1 and 3. For AVR-PikD/OsHIPP19 n = 16 in replicates 1 and 3 and n = 24 in replicate 2. Data points are shown with shapes indicating the replicate. Different letters indicate statistically significant differences based on a pairwise Wilcoxon test (α = 0.05).

The lack of interaction between AVR-PikC and AVR-PikD with FL OsHPP09, OsHPP10, OsHPP11 and OsHIPP21 was validated through split-luciferase complementation assays (**Fig. 5b**). AVR-Pia showed an interaction with OsHIPP19 similar to that observed for AVR-PikD, and a weaker interaction with OsHIPP20 (**Fig. 5b**). Co-expression of Pwl2 with OsHPP09, OsHPP10, OsHPP11 and OsHIPP21 resulted in very weak luminescence. Protein expression was confirmed by immunoblot (**Supplementary. Fig. 15**).

Together, our findings indicate that the HMA domains bound by AVR-Pia are not targeted by AVR-Pik and Pwl2, suggesting that despite sharing a conserved MAX effector fold, these effectors target different H(I)PPs.

### The crystal structure of the AVR-Pia/OsHPP09-HMA complex reveals interface contacts underpinning high affinity binding

To understand the structural basis of the interaction between AVR-Pia and OsHPP09-HMA, we purified and crystallised the effector/HMA complex. Needle-like crystals were obtained in commercial screens and subsequent optimisation of conditions resulted in crystals suitable for diffraction studies. X-ray diffraction data were collected at the European Synchrotron Radiation Facility (ESRF) to a resolution of 1.65 Å. The structure was solved by molecular replacement, refined and validated as described in the Materials and Methods. Data collection, processing and refinement statistics are presented in **Supplementary Table 4**. The final refined model has been deposited in the Protein Data Bank (PDB accession code 9RSV).

As expected, OsHPP09-HMA adopts the well-characterised HMA domain fold, consisting of a four-stranded antiparallel β-sheet and two ⍺-helices arranged with β⍺ββ⍺β topology (**Fig. 6**). Structures of AVR-Pia have previously been determined by NMR spectroscopy^4,52^ (PDB accession codes 2MYW, 2N37) and X-ray crystallography^48^ (PDB accession code 6Q76; complex with Pikp-1 HMA ID). The structure of AVR-Pia determined here is highly similar (RMSD of 0.357 Å to 6Q76, 0.951 Å to 2MYW, and 1.690 Å to 2N37) and adopts the six-stranded β-sandwich fold characteristic of the MAX effectors (**Fig. 6**)^4,6^.

**Fig. 6.**
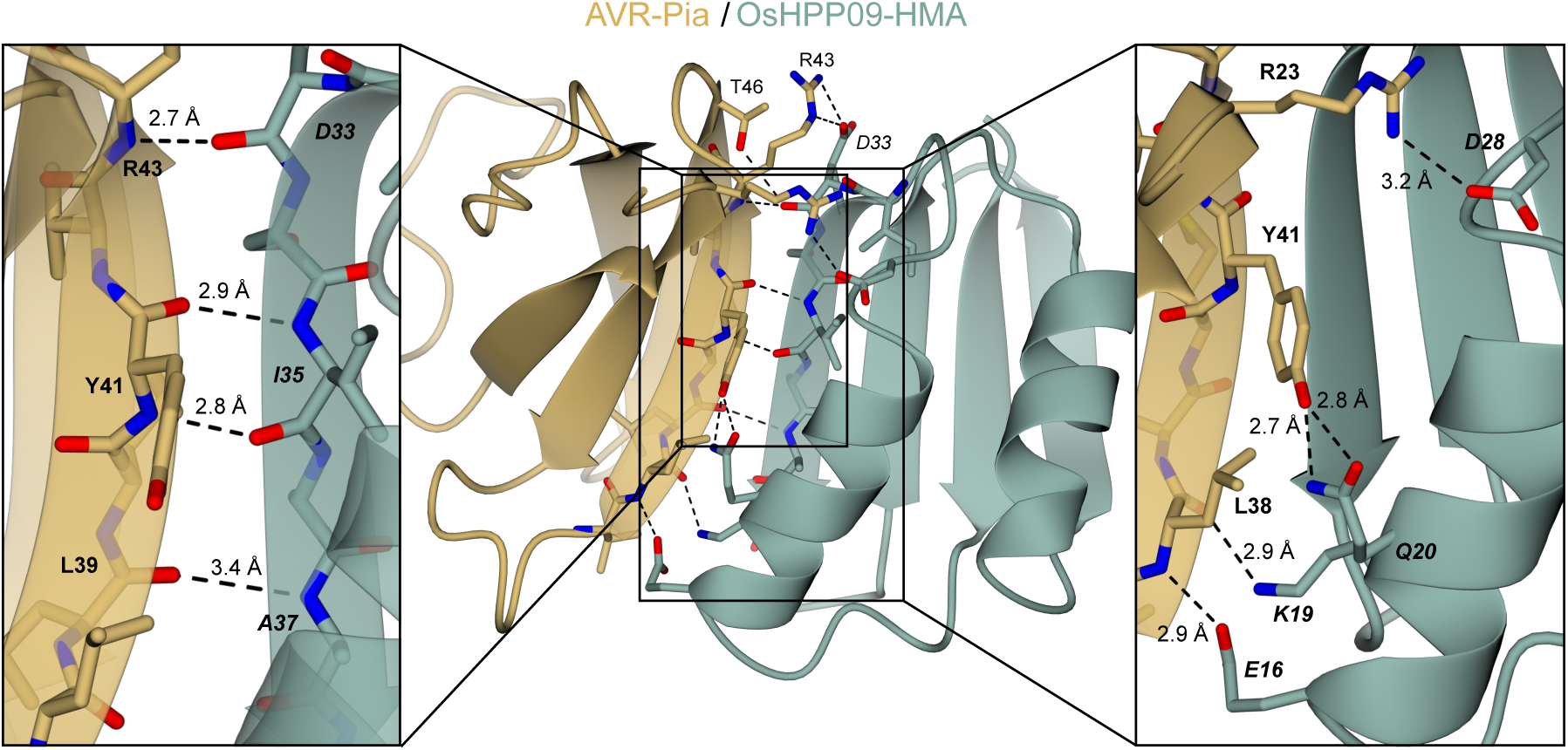
Crystal structure of the HMA domain of OsHPP09 in complex with AVR-Pia. The structures of OsHPP09-HMA and AVR-Pia are represented as teal and gold ribbons, respectively. The amino acids comprising β2 of AVR-Pia and β2 of the HMA domain are represented as cylinders. Side chains involved in intermolecular hydrogen bonds/salt bridges are also represented as cylinders. Hydrogen bonds are represented by black dashed lines labelled with the bond length. Left insert shows main chain hydrogen bonds between AVR-Pia-β2 and OsHPP09-β2. Right insert shows intermolecular contacts between side chains of residues in OsHPP09-⍺1 and AVR-Pia.

Globally, the AVR-Pia/OsHPP09-HMA complex resembles the previously determined structures of AVR-Pia/OsPikp-1-HMA (PDB 6Q76) and AVR1-CO39/OsRGA5-HMA (PDB 5ZNG) (**Supplementary Fig. 16**). The AVR-Pia/OsHPP09-HMA interface area is 521.7Å^2^, fractionally larger than AVR-Pia/OsPikp-HMA (460.7 Å^2^) and AVR1-CO39/OsRGA5-HMA (492.8 Å^2^) (**Supplementary Table 5**). In all three complexes, β2 of the MAX effector is aligned with β2 of the HMA domain, forming an antiparallel β-sheet comprising β6, β1, and β2 of the MAX effector and the four-stranded β-sheet of the HMA domain (**Supplementary Fig. 16a**), with hydrogen bonding between adjacent β-strands (**Supplementary Fig. 16b**).

In contrast to AVR-Pia/OsPikp-HMA and AVR1-CO39/OsRGA5-HMA, where the interface is dominated by main chain hydrogen bonding, in OsHPP09-HMA/AVR-Pia, the side chains of multiple residues in ⍺1 of the HMA domain form intermolecular hydrogen bonds with the effector. The side chains of OsHPP09^E16^ and OsHPP09^K19^ form hydrogen bonds with the peptide backbone of AVR-Pia^L38^ (**Supplementary Fig. 16c**). OsPikp-1^R203^ is in the corresponding position to OsHPP09^K19^, however it is OsPikp-1^R226^, in the β2-β3 loop, which forms an analogous hydrogen bond with AVR-Pia^L38^ (**Supplementary Fig. 16c**). OsRGA5-HMA has a basic residue (R1012) at the corresponding position to OsHPP09^K19^, and the side chain of OsRGA5-HMA^R1012^ is orientated towards the AVR1-CO39^D35^ side chain, however the distance between them (4.1 Å) is greater than the 4 Å cutoff typically used when considering salt bridge interactions (**Supplementary Fig. 16c**). At the start of β2, the OsHPP09^D33^ side chain forms salt bridge interactions with the AVR-Pia^R43^ side chain (**Supplementary Fig. 16d**). This aspartate residue is conserved in both OsPikp-HMA (D217), where it similarly forms a salt bridge with AVR-Pia^R43^, and OsRGA5-HMA (D1026), where it forms an indirect, water-mediated contact with the backbone of AVR1-CO39^T41^ (**Supplementary Fig. 16d**). The OsHPP09^Q20^ side chain forms hydrogen bonds with the side chain hydroxyl of AVR-Pia^Y41^ (**Supplementary Fig. 16e**). The serine in the corresponding position in OsPikp-HMA (S204) forms a water-mediated contact with the side chain hydroxyl of AVR-Pia^Y41^ (**Supplementary Fig. 16e**). Finally, towards the C-terminal end of OsHPP09-⍺1, the side chain of OsHPP09^D28^ forms a salt bridge with AVR-Pia^R23^ (**Supplementary Fig. 16f**). By contrast, the corresponding amino acid in OsPikp-1, OsPikp-1^S212^, forms a hydrogen bond with the side chain hydroxyl of AVR-Pia^Y85^ (**Supplementary Fig. 16f**). Overall, the increased number of intermolecular hydrogen bonds in the AVR-Pia/OsHPP09-HMA structure compared to the structure of AVR1-CO39/OsRGA5-HMA likely explains the difference in binding affinities.

To validate the interface observed in the crystal structure, NMR titration with ^1^H-^15^N-HSQC NMR 2D spectra of ^15^N-labelled AVR-Pia alone and in the presence of unlabelled OsHPP09-HMA were performed. HSQC spectra for ^15^N-labelled AVR-Pia had previously been collected, and the resonances of amide cross-peaks had been assigned^4^. Binding of AVR-Pia to OsHPP09-HMA alters the chemical environment of the amino acids located at the binding interface, resulting in a change in the chemical shift observed for those amino acids in the ^15^N HSQC spectra. The ^15^N HSQC spectra of the complex AVR-Pia with unlabelled OsHPP09-HMA were reassigned using ^15^N,^13^C-AVR-Pia and two- and three-dimensional NMR experiments **(Supplementary Fig. 17).** Depending on the exchange rate constant (kex) between the bound and unbound states and on the frequency difference (Δω) between the corresponding resonances in these two states, different NMR exchange regimes occur. NMR titration showed that the AVR-Pia:OsHPP09-HMA complex was in slow exchange with kex << Δω since separate resonances appeared for individual species **(Supplementary Fig. 18)**. An NMR slow exchange regime indicates a relatively high affinity (<0.1 µM) between the two proteins. The chemical shift perturbations of AVR-Pia in the absence and presence of OsHPP09-HMA were calculated, and significant changes in chemical shift (> 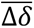 + 1σ) were observed for multiple residues in β2 and β3 of AVR-Pia, consistent with the interface in the crystal structure **(Supplementary Fig. 19)**.

### Replacement of OsHPP09^E16^ with alanine or arginine disrupts AVR-Pia binding

We aimed to identify point mutations in OsHPP09-HMA that interfere with AVR-Pia binding. First, we selected residues whose side chains contribute intermolecular hydrogen bonds or salt bridges and mutated these to alanine (E16A, K19A, Q20A, D28A, D33A) and, for acidic residues, to arginine (E16R, D28R, D33R). D33 is conserved in OsHPP09, OsHPP10, OsHPP11, and in the HMA IDs of Pikp-1 (D217) and RGA5 (D1026) (**Supplementary Fig. 16**D). By contrast, the corresponding residue in Pikm-HMA, which does not bind AVR-Pia^48^, is histidine. We therefore also mutated D33 to histidine. Second, we used FoldX^53^ to predict the effect of mutating each interface residue in OsHPP09-HMA to all other amino acids. Based on the calculated difference of interaction free energy (ΔΔG) upon mutation, we made three further mutations: K19W, I35K, and A37W.

Introduction of the point mutations E16A, E16R, D28A, D28R and I35K in OsHPP09 resulted in a complete loss of interaction with AVR-Pia in Y2H experiments (**Fig. 7a**). The remaining mutations (i.e., K19A, K19W, D33A, D33H, D33R, A37W) with the exception of Q20A, led to reduced yeast growth on selective media suggesting a weakening of the interaction compared to the wildtype OsHPP09/AVR-Pia pair (**Fig. 7a**). Immunoblot analysis revealed that the K19A, K19W and I35K mutants exhibited lower protein expression levels, while all other mutant constructs showed similar expression to wildtype OsHPP09 (**Supplementary Fig. 3d**).

**Fig. 7.**
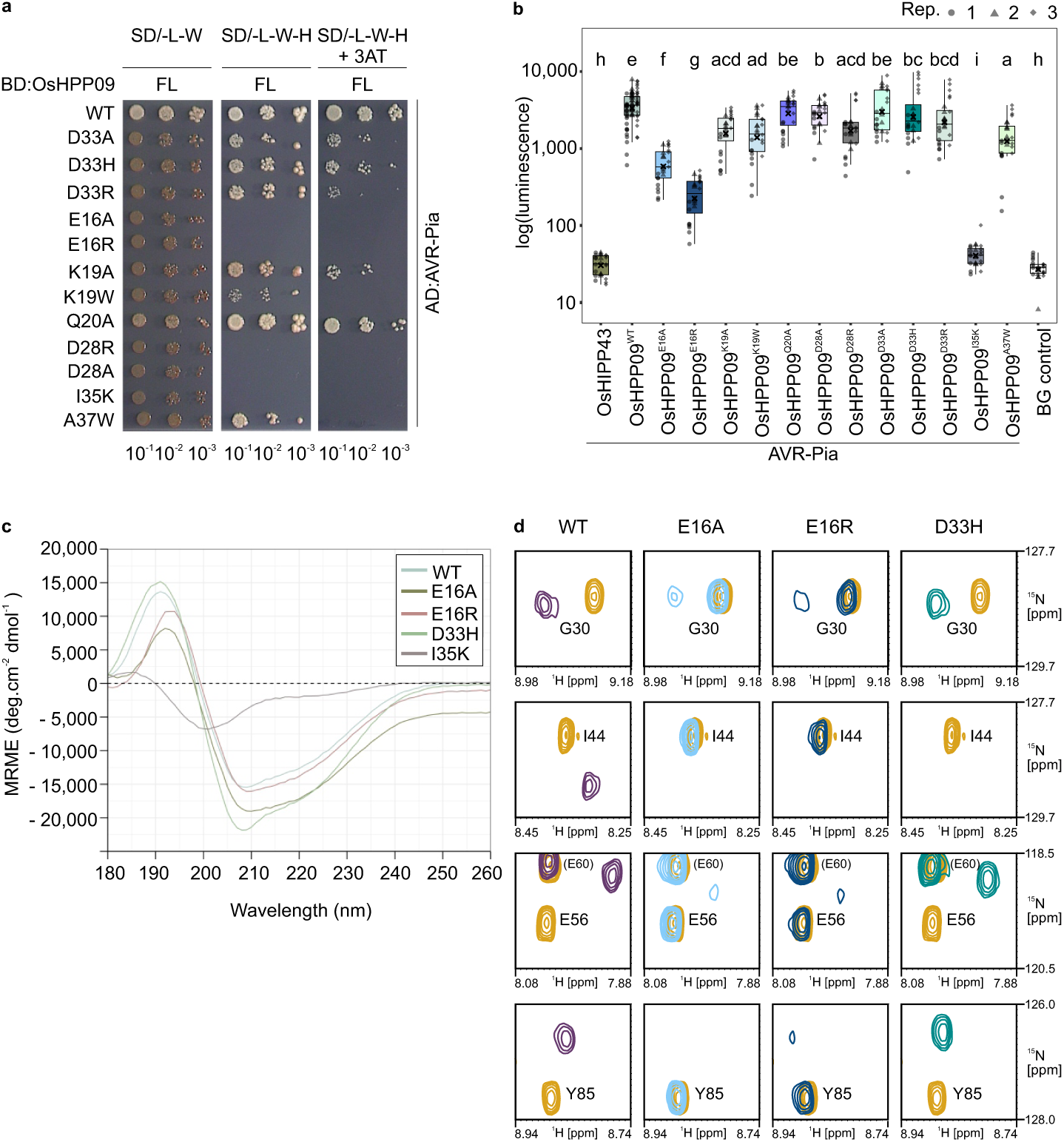
Mutation of OsHPP09 E16 reduces AVR-Pia binding. **a** Y2H interaction analysis between AVR-Pia (ΔSP) and full-length (FL) OsHPP09 carrying single point mutations. OsHPP09^WT^/AVR-Pia served as positive control. Serial dilutions of diploid yeast were spotted onto synthetic defined (SD) media to monitor growth (SD/-LW) or to assess protein-protein interactions (SD/-LWH and SD/-LWH supplemented with 5 mM 3AT). Photos were taken after 7 days of incubation. AD, activating domain; BD, binding domain. **b** Split luciferase complementation assays of AVR-Pia (ΔSP; fused to the N-terminal part of luciferase; 3xHA:NLuc:AVR-Pia) with FL OsHPP09 carrying indicated mutations (fused to the C-terminal part of luciferase: 3xFlag:CLuc:OsHPP09). Co-expression of AVR-Pia with OsHPP09 served as a positive control, co-expression of OsHIPP43 with AVR-Pia as a negative control. Leaves infiltrated with P19 alone served as background (BG) control. Box plots show the median (line), mean (cross) and upper/lower quartiles (box limits), whiskers extend to the most extreme data points within 1.5x the interquartile rang, with outliers plotted individually. Three independent replicates were performed, with n = 8 per combination per replicate. Data points are shown with shapes corresponding to the replicate. Different letters indicate statistically significant differences based on a pairwise Wilcoxon test (α = 0.05). **c** Circular dichroism (CD) spectra obtained for WT OsHPP09-HMA (pale blue) and variants carrying E16A, E16R, I35K or D33H mutations. Negative peaks at 222 nm/208 nm and a positive peak at 193 nm indicate ⍺-helices while a negative peak at 218 nm and positive peak at 195 nm indicate β-sheets, consistent with the β⍺ββ⍺β HMA domain topology. Low ellipticity above 210 nm and a negative peak at 200 nm indicates disorder. **d** Overlays of cross-peaks from HSQC spectra corresponding to residues G30, I44, E56 and Y85 of ^15^N-labelled AVR-Pia alone (orange) and in the presence of either WT OsHPP09-HMA or variants carrying E16A, E16R or D33H mutations (blue/green/purple). Overlays of the full spectra are provided in **Supplementary** Figures 21-23.

To assess the effect of the mutations in the context of the full-length protein in planta, we performed split-luciferase complementation assays. The E16A and E16R mutants showed significantly reduced interaction with AVR-Pia, while the I35K mutant completely lost its ability to interact with AVR-Pia, consistent with the Y2H results (**Fig. 7b**). In contrast, the D28A and D28R mutations, which abolished interaction in Y2H assays, had only a minor effect on the interaction with AVR-Pia in planta, similar to the other tested mutants (**Fig. 7b**). Immunoblot analysis demonstrated that all proteins were expressed at comparable levels in planta (**Supplementary Fig. 20**).

Next, we aimed to establish whether the E16A, E16R, D33H and I35K mutations affect the structure of the HMA domain. Mutated HMA domains were produced in *E. coli*, purified to homogeneity, and molecular weights were confirmed by intact mass spectrometry. Circular dichroism (CD) spectra for OsHPP09^E16A^-HMA, OsHPP09^E16R^-HMA and OsHPP09^D33H^-HMA were comparable to the spectra for wildtype OsHPP09-HMA with features characteristic of ⍺-helices (negative peaks around 222 nm and 208 nm, positive peak around 193 nm) and β-sheets (negative peak around 218 nm, positive peak around 195 nm) consistent with the HMA domain fold (**Fig. 7c**). By contrast, OsHPP09^I35K^-HMA showed features of disorder (low ellipticity above 210 nm and a negative peak around 200 nm) (**Fig. 7c**), suggesting that the lack of AVR-Pia binding to the I35K mutant is likely due to disruption of the HMA domain fold.

We obtained ^1^H-^15^N-HSQC NMR spectra of ^15^N-AVR-Pia in the presence of (unlabelled) HMA domains of OsHPP09^E16A^, OsHPP09^E16R^, and OsHPP09^D33H^. The spectra of ^15^N-AVR-Pia with OsHPP09^D33H^-HMA was highly similar to the spectra of ^15^N-labelled AVR-Pia with WT OsHPP09-HMA (**Fig. 7d**, **Supplementary Fig. 21**), although the exchange regime changes at certain cross-peaks, suggesting lower binding affinity. By contrast, the spectra of ^15^N-AVR-Pia with OsHPP09^E16A^ or OsHPP09^E16R^ showed no chemical shift perturbation and largely superimposed with the spectra of ^15^N-AVR-Pia alone (**Fig. 7d**, **Supplementary Fig. 22**, **Supplementary Fig. 23**). Taken together, these results demonstrate that E16A and E16R mutations block AVR-Pia binding without disrupting the HMA fold, while the I35K mutation interferes with correct folding of the HMA domain.

## Discussion

In the present study, we found that the *M. oryzae* MAX effector AVR-Pia binds to the HMA domains of four rice H(I)PPs: OsHPP09, OsHPP10, OsHPP11, and OsHIPP21. Neither AVR-Pik nor Pwl2 interacts with these H(I)PPs, demonstrating that AVR-Pia binds a distinct group of H(I)PPs to those previously identified as MAX effector targets. Indeed, AVR-Pik interacts with multiple H(I)PPs within a large phylogenetic clade (referred to as “clade A”)^27^, while Pwl2 interacts specifically with OsHIPP43 and its homologs from other grass species^9^.

Strikingly, the interactions between each of these three MAX effectors and HMA domains involve distinct interfaces. In the crystal structures of AVR-Pia/OsHPP09-HMA and Pwl2/OsHIPP43-HMA, β2 of the core MAX fold and β2 of the HMA domain are aligned in an antiparallel orientation^9^, forming an antiparallel β-sheet comprising the four β-strands of the HMA domain and β2, β1, and β6 of the MAX fold. When the HMA domains are superposed, the relative positions of β2 of the MAX fold of Pwl2 and AVR-Pia are very similar (**Supplementary Fig. 24**). However, the other β-strands of Pwl2 are shifted compared to AVR-Pia, enabling residues in β3, β4, and the β3-β4 loop of the MAX fold to contribute to the HMA-binding interface. By contrast, in the OsHIPP19-HMA/AVR-PikF complex, an antiparallel β-sheet forms from the four β-strands of the HMA domain and β3, β4 and β5 of the MAX fold, mediated by hydrogen bonding between β3 of AVR-PikF and β4 of HIPP19-HMA^8^. Furthermore, while AVR-Pia consists solely of the core MAX fold, AVR-Pik has an unstructured extension at the N-terminus, while in Pwl2 the characteristic MAX fold is followed by an ⍺-helix and a C-terminal unstructured region, and these extensions are involved in HMA binding (**Supplementary Fig. 24**). Pwl2^E89^, in the ⍺-helix, forms a hydrogen bond with OsHIPP43-HMA^K51^, while the C-terminal unstructured segment of Pwl2 extends across the HMA domain, forming an extensive interface involving both hydrogen bonds and π-stacking interactions. Similarly, the N-terminal extension of AVR-Pik forms multiple hydrogen bonds with residues in β3, β2, and the β2-β3 loop of OsHIPP19-HMA. The interface in the AVR-Pia/OsHPP09-HMA complex is smaller than in AVR-PikF/OsHIPP19-HMA and Pwl2/OsHIPP43-HMA (521.6 Å^2^ compared to 1044.3 Å^2^ and 1965.2 Å^2^, respectively), and is formed almost exclusively by residues in β2 of the core MAX fold.

It is highly interesting that at least three MAX effectors have evolved to bind HMA domains in different ways. This highlights how structural variation within this effector family can underpin different interactions with host targets. It is unknown whether the interactions of AVR-Pia, AVR-Pik, and Pwl2 with OsH(I)PPs lead to similar outcomes (redundancy) or if each binding event has a distinct and specific impact on plant susceptibility and/or fungal virulence.

While AVR-Pia, AVR-Pik, and Pwl2 all target OsH(I)PPs, the MAX effector AvrPiz-t has been reported to interact with a range of host proteins^54,55^. From a structural perspective, it is unclear how a single MAX effector binds to such diverse targets. However, this suggests that MAX effectors may target many different host proteins and processes during infection.

AVR-Pia is recognised by the paired rice NLR immune receptors OsRGA5 and OsRGA4 through direct binding of the effector to the non-canonical HMA domain integrated into OsRGA5^11^. Such non-canonical domains are hypothesised to have originated in effector host targets and were incorporated into NLR proteins to function as effector “sensors”^15^. Consistent with this, the crystal structure of AVR-Pia/OsHPP09 revealed that AVR-Pia binds OsHPP09-HMA and OsRGA5-HMA through similar interfaces^13^. Interestingly, the F24S and T46N polymorphisms of AVR-Pia-H3 interfered with binding to both OsRGA5-HMA and the candidate OsH(I)PP targets, in contrast to naturally-occurring polymorphisms in AVR-Pik which prevent binding to the HMA ID of Pikp-1 without disrupting interaction with the HMA domains of OsH(I)PP19 and OsH(I)PP20^8,27^.

AVR1-CO39 is also recognised by direct binding to the HMA ID of OsRGA5. However, AVR1-CO39 did not interact with the same OsH(I)PPs as AVR-Pia. AVR1-CO39 also did not interact with the “clade A” OsH(I)PPs bound by AVR-PikD^27^. AVR1-CO39 is broadly absent from rice-infecting isolates of *M. oryzae*^7,56,57^ and its loss has potentially contributed to a host shift from foxtail millet (*Setaria* sp.) to rice. It is therefore possible that AVR1-CO39 has evolved to target H(I)PPs from other grass species, and does not interact with those present in rice.

Interestingly, phylogenetic analysis based on HMA domains clusters OsRGA5-HMA with the H(I)PPs targeted by AVR-Pik and the HMA ID of OsPik-1, and not with OsHPP09, OsHPP10, OsHPP11, and OsHIPP21^27^. This suggests that the RGA5-ID did not arise through duplication and integration of these AVR-Pia candidate targets, but instead from another H(I)PP. We observed interaction between AVR-Pia and FL OsHIPP19, but not with the HMA domain alone, in low stringency Y2H experiments and split-luciferase assays. This suggests that the C-terminal part of OsHIPP19 contributes to AVR-Pia binding and merits further study.

A previous study reported that AVR-Pia binds OsRGA5-HMA with a *K*_D_ of 7.8 μM^13^. This is comparable to the binding affinity of AVR-Pia to OsHPP10-HMA, but at least one order of magnitude weaker than the AVR-Pia/OsHPP09-HMA interaction. This higher affinity can be explained by the number of intermolecular hydrogen bonds and salt bridges at the AVR-Pia/OsHPP09 interface; available structures of AVR1-CO39/OsRGA5-HMA and AVR-Pia/OsPikp-HMA, combined with modelling approaches, suggest that the AVR-Pia/OsRGA5-HMA interface likely involves fewer intermolecular contacts. A similar pattern was reported for AVR-Pik/HMA interactions; surface plasmon resonance experiments showed that AVR-PikD interacts with the HMA domain of OsHIPP19 with higher affinity than the HMA ID of OsPik-1^8^. It is important to note that these binding affinities are determined in vitro using purified HMA domains. These experiments therefore lack the biological context of the FL protein or the plant cell environment.

H(I)PPs, which form extensive families in plants^19,25,26^, have been reported to be involved in intracellular metal transport and homeostasis, responses to cold and drought stress, and heavy metal tolerance. In addition to the interactions between *M. oryzae* MAX effectors and OsH(I)PPs, there are multiple examples of plant H(I)PPs targeted by proteins from diverse pests and pathogens. For example, the putative effector RsMf8HN from the necrotrophic fungus *Rhizoctonia solani* interacts with OsHIPP28^38^, while a HIPP from sugarcane (ScPi21) is a target for the *Fusarium sacchari* effector Fs00367^35^. CsHIPP03 was identified by Y2H screening as an interactor of SDE34, an effector secreted by the phloem-colonising bacteria *Candidatus* Liberibacter asiaticus^40^. A root-knot nematode (*Meloidogyne graminicola*) effector binds rice OsHPP04 and suppresses the flg22-induced ROS burst^36^. Finally, the potato mop-top virus movement protein interacts with NbHIPP26 to promote systemic spread of the virus^31^. Together, these results suggest that the manipulation of H(I)PPs is required for successful infection by diverse pests and pathogens. However, little is known about the molecular mechanisms through which H(I)PPs affect plant resistance and/or susceptibility.

HMA domains typically coordinate metal ions via two cysteine residues in an MxCxxC motif. Intriguingly, the metal-binding motif is degenerate in all four HMA domains bound by AVR-Pia, with both cysteines replaced by other amino acids (MT<u>D</u>EK<u>T</u> in OsHPP09, MT<u>D</u>DKI in OsHPP10, MS<u>D</u>TK<u>M</u> in OsHPP11, and MT<u>D</u>ER<u>K</u> in OsHIPP21), indicating the absence of metal-binding. Consistent with this, no electron density suggestive of bound metal ions was observed in the OsHPP09-HMA/AVR-Pia crystal structure. Interestingly, OsHPP09^E16^, which is important for AVR-Pia binding, is located in the degenerate metal binding motif. Of the “clade A” H(I)PPs bound by AVR-Pik, some contain an intact MxCxxC motif, while others, such as OsHIPP19 and OsHIPP20, lack one or both cysteine residues. OsHIPP43 has an intact MxCxxC motif, though it is unknown whether it binds metal ions in planta. In cases where HMA domains lack a complete MxCxxC motif, and presumably also the capacity to bind metals, the functions of the H(I)PPs are particularly intriguing. Functional diversification, beyond metal transport and homeostasis, may have been facilitated by the stable core fold of the HMA domain supporting diversification of surface properties to deliver new functions. Protein prenylation can affect subcellular localisation, protein stability, and protein-protein interactions^58^. For some HIPPs, the C-terminal CaaX isoprenylation motif has been shown to drive localisation to plasmodesmata. Further investigation is required to establish the localisation and function of the AVR-Pia interactors in planta.

Pathogens require certain host factors to establish a successful infection, with the underlying genes typically referred to as susceptibility (S) genes. Previous studies have shown that *OsHIPP05* (*Pi21*) and *OsHIPP20* are S genes for blast infection; *oshipp05* and *oshipp20* CRISPR-Cas9 knockout plants exhibited increased blast resistance compared to WT plants ^27,33,34,59–61^. Further, a recent study reported that overexpression of barley (*Hordeum vulgare*) *HvHIPP43* increased susceptibility to *M. oryzae,* suggesting that *HvHIPP43* may also be an S gene^34^. Other plant H(I)PPs are also S genes for different pests and pathogens; *Arabidopsis thaliana* plants carrying loss-of-function mutations in *AtHIPP27* or *AtHMAD* (also known as *AtHIPP40*^29^) exhibited reduced susceptibility to the beet cyst nematode *Heterodera schachtii*^32^ and the bacterial pathogen *Pseudomonas syringae*^41^, respectively. Silencing of *TaHIPP1* in wheat reduced sporulation of the wheat stripe rust fungus *Puccinia striiformis* f. sp. *tritici*^39^. *OsHPP09*, *OsHPP10*, *OsHPP11,* and/or *OsHIPP21* may also be S genes for rice blast disease, and deletion of these genes may increase blast resistance.

Knocking out S genes has the potential to confer durable resistance, but can cause undesirable pleiotropic effects on plant growth and development^62–65^. Base editing offers a more precise alternative, allowing single amino acid changes that minimise disruption of protein function^66^. While the functions of OsHPP09, OsHPP10, OsHPP11, and OsHIPP21, and the consequences of AVR-Pia binding remain unknown, the virulence function of the MAX effector Pwl2 depends on interaction with OsHIPP43^34^. Altering effector targets to block effector binding could compromise the virulence function of effectors without compromising the folding or endogenous function of the host protein. However, it is crucial to verify that such mutations do not interfere with the folding and function of the host targets before commencing time-consuming and costly base editing experiments in plants.

In summary, we have characterised the interactions between AVR-Pia and a novel subset of HMA domain-containing rice proteins. Biophysical and structural analyses guided mutagenesis of these effector targets and identified a mutation that blocks effector binding without compromising structural integrity of the HMA domain. This provides a foundation for targeted modification of these proteins for enhanced disease resistance.

## Methods

### Cloning

The coding sequences (CDS) of *OsH(I)PPs*, including the stop codon, were synthesised by Twist Bioscience. The CDS of *OsHIPP14*, *OsPi21*, *OsHIPP19*, *OsHIPP21,* and *OsHPP09* were codon-optimised to enable synthesis.

For Y2H and split luciferase experiments, synthesised *OsH(I)PP* genes were used as template for amplification with primers containing *attB1* and *attB2* recombination sites (**Supplementary Table S6**) and Phusion High-Fidelity DNA Polymerase (Thermo Fisher Scientific). Resulting PCR products were subsequently introduced into the pDONR207 vector (Invitrogen) via Gateway® BP recombination (Thermo Fisher Scientific). Entry constructs for the MAX effectors dSP-AVR-Pia^15^, dSP-AVR1-CO39^12^, dSP-AVR-PikD^47^ and dSP-Pwl2^13^ were previously published. The dSP-AVR-PikC fragment was amplified from pCambia1300_dSP-AVR-PikC^45^ and dSP-MAX58 from a synthesised gene, both using primers containing *attB1* and *attB2* recombination sites (Supplementary Table S6) and Phusion High-Fidelity DNA Polymerase (Thermo Fisher Scientific). Resulting PCR products were cloned into the pDONR207 vector (Invitrogen) via Gateway® BP recombination (Thermo Fisher Scientific).

For recombinant protein production in *E. coli,* synthesised genes were used as templates for PCR amplification of HMA domains (**Supplementary Table S1**) with primers containing *Bpi*I restriction sites (**Supplementary Table S6**) using CloneAmp™ HiFi PCR Premix (Clontech Laboratories). Resulting PCR products were introduced into pICH41308 (Addgene #47998^67^) by Golden Gate assembly^68^ to generate level 0 CDS modules. The CDS of OsHIPP21-HMA was codon-optimised for *E. coli* expression and supplied with appropriate overhangs and restriction sites for direct assembly into the vector pICH41308 without PCR amplification. Level 0 CDS modules were used in Golden Gate reactions with the pOPIN-GG vector pPGN-C (Addgene #174578^69^) and the level 0 module pICSL30015 (Addgene #174582^69^) to generate *E. coli* expression constructs with a cleavable N-terminal 6xHis-MBP tag.

Site-directed mutagenesis of OsHPP09-HMA for Y2H, split luciferase, and recombinant protein production was performed using a QuikChange Site-Directed Mutagenesis Kit (Agilent). Prenylation motif mutations (*OsHIPP14^C187S^*, *OsHIPP21^C133S^*, *OsHIPP39*^C190S^, *OsHIPP41*^C152S^) in the pDONR constructs, and mutations in AVR-Pia to give *AVR-Pia-H3*, were introduced by site-directed mutagenesis PCR using Phusion High-Fidelity DNA Polymerase (Thermo Fisher Scientific) and mutation-specific primers listed in **Supplementary Table S6**.

Gateway® LR recombination using LR Clonase® II mix (Thermo Fisher Scientific) was used to transfer *OsH(I)PPs* and *RGA5_Cter_*^11^ (883-1116 aa) into Gateway®-compatible destination vectors, including a modified Y2H vector pGBKT7-GW^70^ and the plant expression vector pBIN-35S-3Flag-Cluc-GWY (Deslandes et al., unpublished). Similarly, MAX effector pDONR constructs were recombined into a modified pGADT7-GW^70^ and pBIN-35S-3HA-Nluc-GWY (Deslandes et al., unpublished) destination vectors. The pBIN-35S-3HA-Nluc-GWY and pBIN-35S-3Flag-Cluc-GWY vectors were generated by ligation of a 3HA-Nluc-FrameB or 3Flag-Cluc-FrameB fragment, respectively, into a XhoI/XbaI digested pBIN-35S vector.

Previously published expression constructs for AVR1-CO39, AVR-Pia, and AVR-Pia-H3^4,12,13^ were used for protein production in the present study.

A complete list of plasmids used in this study is provided in **Supplementary Table S7.**

### Yeast two-hybrid

The yeast strain Y2HGold (mating type a; Takara Bio) was transformed with the bait construct expressed from the pGBKT7 vector, which contains the GAL4-BD and the *TRP1* marker gene. These constructs included previously described HMA domains ^49^, as well as newly generated constructs: OsHIPP21-HMA, OsH(I)PP^WT^ FL and OsH(I)PP^C/S^ FL. The yeast strain Y187 (mating type α; Takara Bio) was transformed with the prey construct (MAX effectors) expressed from the pGADT7 vector, which carries the GAL4-AD and the *LEU2* marker gene. Yeast transformation was performed using a lithium acetate (LiAc)-based protocol according to the Yeastmaker Yeast Transformation System (Takara Bio). Transformed yeast cells were plated on synthetic defined (SD) media lacking either tryptophan (-W) or leucine (-L) and incubated at 30 °C for 3 days.

Mating of the transformed yeast strains followed the Matchmaker Gold Yeast Two-Hybrid System User Manual (Takara Bio). Diploid yeast cells were selected on SD/-L-W plates (SD medium lacking leucine and tryptophan) after incubation at 30 °C for 2 days. For Y2H assays, diploid yeast cells were inoculated in SD/-L-W medium, grown for 24 hours at 28°C with shaking at 100 rpm, then pelleted and washed twice with sterile water. Serial 10-fold dilutions (10⁻¹, 10⁻², and 10⁻³) were spotted onto SD selection plates: SD/-L-W (growth control), SD/-L-W-H (lacking leucine, tryptophan, and histidine), SD/-L-W-H supplemented with varying concentrations of 3-amino-1,2,4-triazole (3AT). After 7 days incubation at 30 °C, plates were photographed using the presentation station UF-130DX (Samsung).

### Yeast two-hybrid cDNA library screening

The Y2H cDNA library derived from the rice cultivar CO39 and the BD:dSP-AVR-Pia construct, have been previously described^11,47^. Library screening followed the Matchmaker Gold Y2H system protocol (Takara Bio). Mating was performed between the Y2H Gold strain harbouring GAL4-BD:AVR-Pia and the Y187 strain carrying the CO39 cDNA library. Diploid yeasts were plated on SD/-L-W-H medium and supplemented with 20 mM 3AT. Colonies that grew were re-streaked onto fresh SD/-L-W-H medium containing 20 mM 3AT to confirm interaction-dependent growth. Yeast colony PCR was used to amplify cDNA inserts from positive clones, and PCR products were sequenced. Sequences were aligned to the cDNAs of two publicly available annotations of the *Oryza sativa* cv. Nipponbare reference genome (IRGSP-1.0^71^ and MSU7.0)^72^ using BLAST to identify the corresponding genes. Only genes for which the cDNA inserts (or fragments of cDNAs) were cloned in-frame with the GAL4-AD were retained as candidate AVR-Pia targets.

### Agrobacterium-mediated transient expression in *N. benthamiana*

*Agrobacterium tumefaciens* strains AGL1 (carrying split-luciferase constructs) and GV3101-pMP90 (carrying 35S::P19) were grown overnight at 28°C in 5 mL LB medium supplemented with appropriate antibiotics. Overnight cultures were pelleted at 4,000 x *g* for 10 minutes, and pellets resuspended in 2 mL infiltration buffer (10 mM MgCl₂, 10 mM MES pH 5.6, 150 µM acetosyringone). Bacterial suspensions were incubated in the dark for 2 hours. The optical density at 600 nm (OD_600_) was adjusted to 0.05 for 35S::P19 and 0.1 for OsH(I)PPs and MAX effectors, respectively. In all infiltration mixes, 35S::P19 was co-infiltrated to suppress gene silencing. Agrobacterial suspensions were mixed and infiltrated into leaves of 5–6-week-old *N. benthamiana* plants using a needleless syringe. Leaf discs were harvested 2 days post-infiltration for luminescence measurements or protein extraction. *N. benthamiana* plants were grown in a growth chamber on soil under a 16-hour light / 8-hour dark cycle, at 20 °C and 65 - 75% relative humidity.

### Split luciferase assay

For luminescence measurements, 4 mm leaf discs (collected using a Miltex Biopsy Punch (Delta Microscopie)) were placed into wells of a white 96-well plate containing water. The water was removed and replaced with a 1 mM Xenolight D-luciferin-K⁺ salt bioluminescent substrate solution (Perkin Elmer). Luminescence was quantified using a Spark® microplate reader (Tecan) over three cycles of 10 minutes each. For all experiments, data from the third cycle were used for analysis. Each interaction was tested with eight technical replicates (eight leaf discs per plate from four individual plants per biological replicate, unless otherwise indicated in the figure legend). The luminescence assay was performed in three independent experiments. Statistical analyses from three independent replicates (24 leaf discs in total, unless otherwise indicated in the figure legend) and data visualisation were done using R (version 4.4.1). Multiple comparisons were performed using pairwise Wilcoxon tests followed by a Benjamini-Hochberg p-value adjustment with the stats package. Boxplots were generated using the ggplot2 package.

### Protein extraction and Western blotting

Protein extraction from yeast was performed using a post-alkaline extraction protocol, as described by Kushnirov^73^. For BD-OsHIPP14/19/20/39/41 and AD-MAX, yeast crude extracts were resuspended in 50 µL of 1x LDS loading buffer (1x Bolt™ LDS sample buffer (Thermo Fisher Scientific), 1x NuPAGE™ sample reducing agent (Thermo Fisher Scientific)), boiled at 95°C for 10 minutes, and loaded onto a 10% NuPAGE™ Bis-Tris gels (Invitrogen). For BD-OsHPP09/10/11 and OsHIPP21, yeast crude extracts were resuspended in 50 µL of 1x SDS-Tricine buffer (1x SDS-Tricine buffer (Thermo Fisher Scientific), 1x NuPAGE™ sample reducing agent (Thermo Fisher Scientific)) and boiled at 85 °C for 2 minutes before loading onto a 10-20% Tricine gel (Novax). Proteins were transferred to iBlot® nitrocellulose membrane (Invitrogen) using the iBlot 2 Gel Transfer Device (program: 20 V 1 min, 23 V 4 min, 25 V, 2 min, Thermo Fisher Scientific), and analysed by immunoblotting.

For protein extraction from *N. benthamiana*, four leaf discs (0.8 mm diameter) were collected, immediately frozen in liquid nitrogen and ground using a tissue homogeniser (Qiagen). The ground plant material was solubilised in 200 µL extraction buffer (20 mM Tris-Hcl pH 7.5, 150 mM NaCl, 1 mM EDTA pH 8.0, 1% Triton X-100, 0.1% SDS, 5 mM DTT, 1x Protease inhibitor cocktail (Sigma-Aldrich), 1x EDTA-free cOmplete protease inhibitor (Roche)). Protein extracts were centrifuged at 18,000 x *g* for 15 minutes at 4 °C, and the resulting supernatants retained. 2x LDS buffer (Bolt™ LDS sample buffer (Thermo Fisher Scientific) and NuPAGE™ sample reducing agent (Thermo Fisher Scientific)) was added and samples were denatured at 70 °C for 10 minutes. Proteins were resolved by electrophoresis on NuPAGE™ 4-12% Bis-Tris gels (Invitrogen). Proteins were transferred to iBlot® nitrocellulose membrane (Invitrogen) as described above and analysed by immunoblotting.

For immunodetection of proteins, rabbit anti-GAL4-BD (sc-577, Santa Cruz Biotechnology, 1:2,000), goat anti-rabbit IgG-HRP (1:20,000, Sigma-Aldrich, A0545), mouse anti-Flag M2-HRP (1:60,000, Sigma-Aldrich) and rat anti-HA-HRP (clone 3F10, 1:1,1000, Roche) were used. Chemiluminescence was detected using the Immobilon western kit (Millipore) or the SuperSignal™ West Femto Maximum Sensitivity substrate (Thermo Fisher Scientific) and the Syngene 680X EF GBOX imaging system (Syngene).

### Protein production and purification

#### HMA domains

HMA domains were produced with a cleavable N-terminal 6xHis-MBP tag from *E. coli* BL21 (DE3) or BL21 (DE3) pLysS. 700 mL cell cultures were grown in LB media at 37 °C from a starting OD_600_ of 0.05-0.07 to an OD_600_ of 0.5-0.8. Protein production was induced by addition of IPTG (final concentration 1 mM) and cultures were grown for a further 14-16 hours at 20 °C. Cells were pelleted by centrifugation and the pellets stored at -70 °C. Thawed pellets were resuspended in 50 mM Tris-HCl pH 7, 300 mM NaCl, 30 mM imidazole supplemented with cOmplete EDTA-free protease inhibitor cocktail (Roche). Cells were lysed by sonication and the lysate clarified by centrifugation at 40,000 x *g* at 6 °C for 30 minutes. Subsequent chromatography steps were carried out with an ÄKTA Pure system (Cytiva) at 4 °C. The filtered (0.45 μm) supernatant was applied to a 5 ml HisTrap^TM^ FF column (Cytiva) and, after washing with 50 mM Tris-HCl pH 7, 300 mM NaCl, 30 mM imidazole, bound protein was step-eluted with 50 mM Tris-HCl pH 7, 300 mM NaCl, 500 mM imidazole. The eluate was injected onto a Superdex 75 26/60 size exclusion chromatography column (Cytiva) pre-equilibrated in 50 mM Tris-HCl pH 7, 150 mM NaCl. Fractions containing the fusion protein were incubated with 3C protease overnight at 4 °C. HMA domains were separated from the 6xHis-MBP tag and GST-tagged 3C protease by injecting the protease-treated sample onto a 5 ml HisTrap^TM^ FF column (Cytiva), 5 ml MBPTrap^TM^ HP column (Cytiva), and 5 ml GSTrap^TM^ HP column (Cytiva) connected in tandem and equilibrated in 50 mM Tris-HCl pH 7, 300 mM NaCl, 30 mM imidazole. Pooled fractions containing HMA domains were concentrated to < 8 ml then injected onto a Superdex 75 26/60 size exclusion chromatography column (Cytiva) pre-equilibrated in 50 mM Tris-HCl pH 7, 150 mM NaCl. Pooled fractions containing the purified HMA domains were concentrated. At each stage, fractions containing the HMA domain were determined by SDS-PAGE. Samples were combined with 5X SDS loading buffer (60 mM Tris-HCl pH 6.8, 25% glycerol, 4% SDS, 0.1% bromophenol blue, 10 mM DTT), heated at 95 °C for 5 minutes, then loaded onto a Bolt™ 10 %, Bis-Tris gel (Invitrogen). After migration, gels were stained in InstantBlue® Coomassie protein stain (Abcam). Final protein concentration was determined using a Pierce™ BCA Protein Assay Kit (ThermoFisher Scientific), due to the extreme molar extinction coefficients of the proteins (0 M^-1^cm^-1^ for OsHIPP21-HMA, 1490 M^-1^cm^-1^ for OsHPP-HMA domains) at 280 nm. Proteins were stored at -80 °C.

#### MAX effectors

AVR-Pia, AVR-Pia-H3 and AVR1-CO39 were produced and purified as previously described^4,12,13^. Concentrations of MAX effectors were determined from molar extinction coefficients and absorbance at 280 nm.

### Analytical size exclusion chromatography

Analytical size exclusion chromatography experiments were conducted using a Superdex 75 10/300 GL column (Cytiva) connected to an ÄKTA Pure system (Cytiva) at 4 °C. Running buffer was 50 mM Tris pH 7, 150 mM NaCl, 1 mM DTT for experiments with AVR1-CO39 and 20 mM sodium citrate pH 5.6, 150 mM NaCl, 1 mM DTT for experiments with AVR-Pia and AVR-Pia-H3. To investigate whether OsHPP09-HMA forms a complex with the effectors, the two proteins were combined in a 1:1 molar ratio and incubated on ice for 1 h prior to analysis. Each protein was also analysed alone, at a concentration equivalent to that present in the mixture. For each experiment, 100 μl protein was injected onto the column and eluted at a flow rate of 0.5 ml/min. 500 μl fractions were collected for analysis by SDS-PAGE (as described in the protein purification section above).

### Circular dichroism spectroscopy

Circular dichroism (CD) spectra were obtained using a Chirascan™ CD Spectrometer (Applied Photophysics). Proteins were dialysed into 10 mM sodium phosphate, pH 7.2 using Slide-A-Lyzer™ MINI dialysis devices (Thermo Fisher Scientific). 40 μl protein was used to fill a 0.1 mm cuvette. The sample compartment temperature was 20 °C. Spectra were obtained for wavelengths from 180 nm to 260 nm with step size of 1 nm, bandwidth of 2 nm and time-per-point of 2 s. Ten spectra were acquired for buffer and each sample. The average trace for buffer was subtracted from the average trace for each protein. Units were converted from millidegrees to mean residue molar ellipticity (MRME) using the Chirascan^TM^ software.

### Nuclear magnetic resonance titration

Samples of ^15^N-labelled AVR-Pia (40 μM) alone and complexed to unlabelled OsHPP09-HMA (1:0.5 or 1:1 molar ratio), or OsHPP09-HMA mutants (1:1 molar ratio), were prepared in 20 mM sodium citrate, pH 5.6, 150 mM NaCl, and 1mM DTT, with addition of 10% D_2_0 for the lock and 1 µM DSS as internal reference for the ^1^H dimension and indirectly referenced for the ^15^N dimension^74^. The backbone (Hn, Nh, Cα) and Cß resonances of ^15^N,^13^C-labeled AVR-Pia complexed to OsHPP09-HMA (1:1 ratio) were assigned using three-dimensional (3D) HNCO, HNCA, HN(CO)CACB, HN(CA)CO and HNCACB experiments recorded at 305K on a Bruker Avance 800 MHz spectrometer equipped with a triple resonance (^1^H, ^15^N, ^13^C) z-gradient cryo-probe, as described in detail for AVR-Pia alone in solution^13^. Spectra were processed using Topspin (v. 3.5pl6) and analysed with Cindy in-house software or CCPN^75^ [analysis v 2.5.2].

Chemical shift perturbations (Δ𝛿) were calculated using the equation^76^:

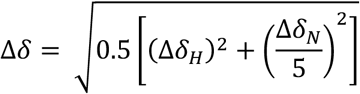

Δ𝛿_𝐻_ and Δ𝛿_𝑁_ are the chemical shift differences measured in the proton dimension and nitrogen dimension, respectively, of 2D [^1^H,^15^N] HSQC spectra recorded for AVR-Pia alone or in the presence of OsHPP09-HMA (1:1 ratio). Δ𝛿 greater than 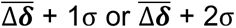 were considered as significant or very significant, respectively, σ being the standard deviation measured on the distribution of chemical-shift perturbations.

### Isothermal titration calorimetry (ITC)

ITC experiments were performed using a MicroCal PEAQ-ITC (Malvern Panalytical). Proteins were dialysed overnight into 20 mM sodium citrate pH 6.0, 150 mM NaCl, 1 mM TCEP. Experiments were carried out at 25 °C with a reference power of 5 μcal/s. The calorimetric cell was filled with purified HMA domain at 18 μM (OsHPP09-HMA), 35 μM (OsHIPP21-HMA and OsHPP11-HMA) or 60 μM (OsHPP10-HMA). This was titrated with AVR-Pia at a concentration 10X greater than that of the HMA domain (180 uM, 350 uM or 600 uM). Each experiment consisted of a single injection of 0.5 μL followed by 18 injections of 2 μL each at 150 s intervals with a stirring speed of 750 rpm. Data were processed with the MicroCal PEAQ-ITC analysis software (Malvern Panalytical). Experiments were carried out in triplicate.

### Crystallisation of OsHPP09-HMA and AVR-Pia

Sitting drop, vapour diffusion crystallisation trials were set up in 96-well Swissci 3 lens crystallisation plates using dragonfly and mosquito dispensing robots (SPT Labtech). 100 nL of protein at 8.6 mg/ml was combined with 100 nL reservoir solution. Needle-like crystals were observed in multiple conditions in the commercial Structure Screen 1 + 2 (HT-96; Molecular Dimensions) and PEGs Suite (NeXtal) screens. Optimisation around these conditions yielded crystals suitable for diffraction studies. The X-ray diffraction data used to solve the structure were obtained from a crystal from 0.1 M MES pH 6.5, 0.2 M ammonium sulphate, 20% (w/v) PEG 4000. The crystal was mounted in a nylon loop and flash-frozen in liquid nitrogen prior to shipping to the European Synchrotron Radiation Facility (ESRF).

### X-ray diffraction data collection and processing, and model refinement

X-ray diffraction data were collected on beamline MASSIF-1 (ID30A-1) at the ESRF. Data were processed with the EDNA autoprocessing pipeline^77^. The scaled, unmerged data file was passed to AIMLESS for data reduction^78,79^ (implemented in CCP4i2^80^). The AVR-Pia/OsHPP09-HMA structure was solved by molecular replacement with PHASER^81^, using a monomer of AVR-Pia (PDB accession 6Q76, chain B) and a monomer of OsHPP09-HMA (modelled with ColabFold^82^). Iterative cycles of manual adjustment, refinement, and validation were carried out using COOT^83^, REFMAC5^84,85^ and MolProbity^86,87^. Analysis of the interface was performed with qtPISA^88^. Interatomic distances were measured and structure superposition/RMSD calculations were performed with PyMOL v2.5.0 Open-Source^89^. Structure figures were prepared using the CCP4 molecular graphics (CCP4mg) software^90^.

### Structural modelling of AVR-Pia/HMA complexes

Structural modelling of AVR-Pia/HMA complexes was carried out with ColabFold v1.5.5 ^82^ (based on AlphaFold2 Multimer v3^91,92^) and the AlphaFold3 web server (alphafoldserver.com; models generated between 2024-05-27 and 2024-07-02)^93^. For models generated with ColabFold, no template information from the PDB was used (template_mode: none) and number of recycles was set to 20 (num_recycles: auto). Five models were generated for each complex and relaxed using AMBER (num_relax: 5). Default settings were otherwise used. For models generated with AlphaFold3, default settings were used. At the time of use, AlphaFold3 did not support excluding template information from the PDB. Five models were generated for each complex using five different seed values. Interface analysis was carried out with qtPISA^88^.

## Acknowledgements

We thank Mark Youles and Mark Banfield for generating and providing the pOPIN-GG vector and tag modules, Laurent Deslandes for providing the split-luciferase vectors, and Florian Veillet for providing the pFV79 construct. We thank Thorsten Langner for providing the Y2H HMA library. We thank Julie Noell for assistance with protein purification, Corinne Lionne and Manon Mallet for helpful discussions about ITC methods, and Christian Roumestand for advice about NMR data. We thank Sebastien Ribeiro, Laura Mathieu and Vincent Chochois for valuable discussions and support with data analysis. This work benefited from access to the NMR and RX facilities of the Integrated Biophysics and Structural Biology Platform (PIBBS) of the CBS. PIBBS is a GIS-IBISA platform and belongs to the French Infrastructure for Integrated Structural Biology (FRISBI), supported by the National Research Agency (ANR-10-INBS-05). We acknowledge the European Synchrotron Radiation Facility (ESRF) for provision of synchrotron radiation facilities. We thank the staff of the ESRF and EMBL Grenoble for assistance in using beamline ID30-A1 under proposal number mx-2409. This work was funded by the HORIZON EUROPE ERC-2019-STG-852482-ii-MAX awarded to Stella Cesari.

## Author contributions

**Josephine H.R. Maidment**: Conceptualization; Data curation; Formal analysis; Investigation; Methodology; Project administration; Resources; Supervision; Validation; Visualization; Writing – original draft; Writing – review & editing. **Svenja C. Saile**: Conceptualization; Data curation; Formal analysis; Investigation; Methodology; Project administration; Resources; Supervision; Validation; Visualization; Writing – original draft; Writing – review & editing. **Aurélien Bocquet**: Investigation; Methodology; Resources; Validation; Writing – review & editing. **Céline Thivolle**: Investigation; Methodology; Resources; Validation; Writing – review & editing. **Léo Bourcet**: Investigation; Methodology; Resources; Validation; Writing – review & editing. **Lisa-Fatimatou Planel**: Investigation; Methodology; Resources; Validation; Writing – review & editing. **Muriel Gelin**: Methodology; Resources; Writing – review & editing. **Thomas Kroj**: Conceptualization; Writing – review & editing. **André Padilla**: Conceptualization; Methodology; Writing – review & editing. **Karine de Guillen**: Conceptualization; Data curation; Formal analysis; Investigation; Methodology; Project administration; Resources; Supervision; Validation; Visualization; Writing – original draft; Writing – review & editing. **Stella Cesari**: Conceptualization; Data curation; Formal analysis; Funding acquisition; Investigation; Methodology; Project administration; Resources; Supervision; Validation; Visualization; Writing – original draft; Writing – review & editing.

## Data availability

The protein structure of the complex between OsHPP09-HMA and AVR-Pia has been deposited in the Protein Data Bank with accession number 9RSV (pdb_00009RSV). X-ray diffraction data are publicly available via the European Synchrotron Radiation Facility ^94^.

