## Supplementary figures S1-S24 and tables S1-S5 for "The *Magnaporthe oryzae* MAX effector AVR-Pia binds a novel group of rice HMA domain-containing proteins"

**This document includes :**

Supplementary Figures S1 – S24
Supplementary Tables S1 – S5
References

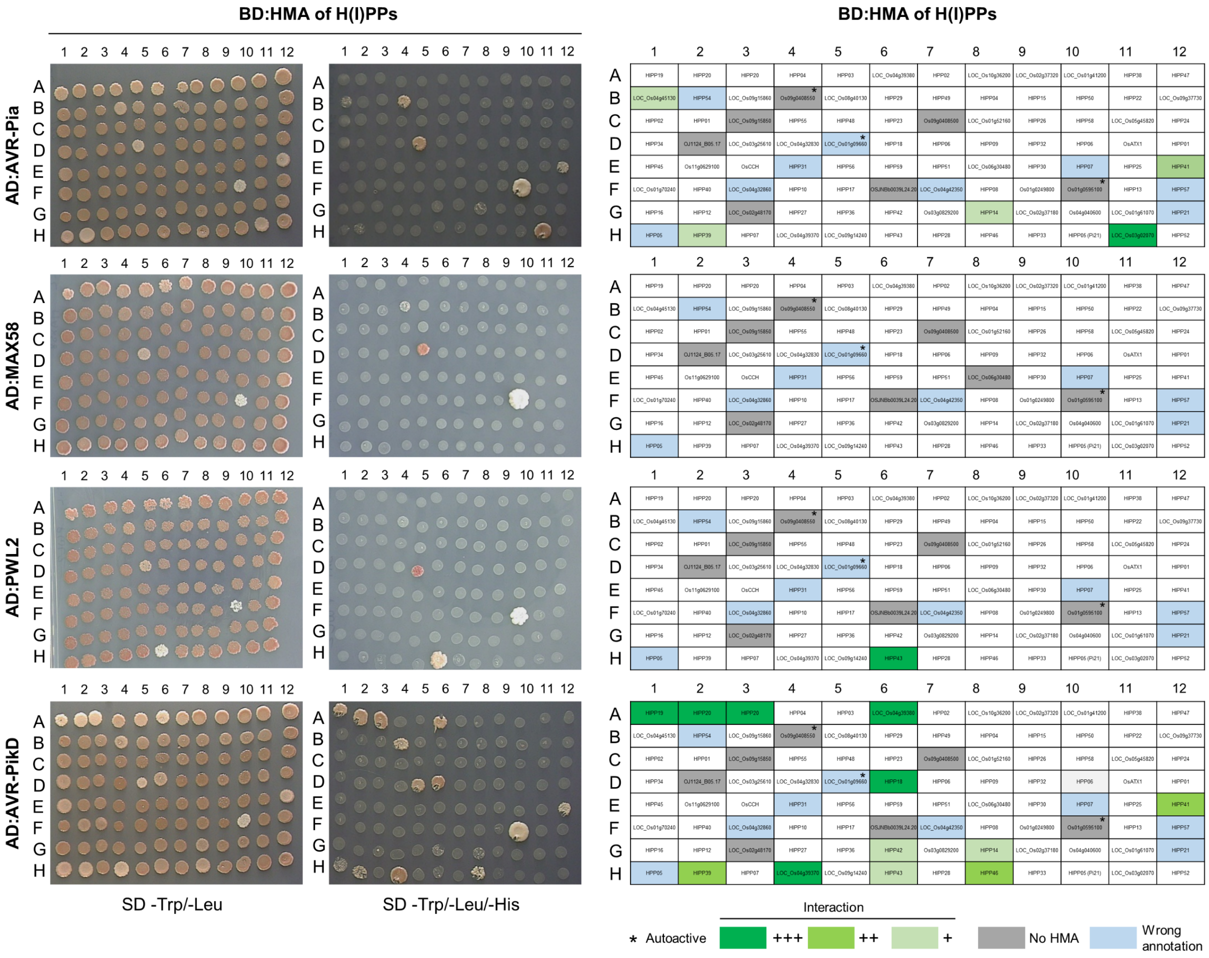


**Supplementary Fig. 1. AVR-Pia interacts with HMA domains from OsH(I)PPs.**

Comprehensive pairwise Y2H analysis using AVR-Pia, MAX58, Pwl2, and AVR-PikD as prey, and a library of HMA domains from OsH(I)PPs as bait[1]. The left panel shows yeast growth on selective media, while the right panel identifies the rice H(I)PP proteins from which the HMA domains were derived. Diploid yeast were spotted onto synthetic defined (SD) media to monitor growth (SD/-LW) or to assess protein-protein interactions (SD/-LWH). Pictures were taken after 7 days of incubation. AD, activating domain; BD, binding domain; *, Autoactive BD construct leading to yeast growth regardless of the presence or identity of the AD construct; +++, strong yeast growth; ++, normal yeast growth, +, weak yeast growth; No HMA, proteins for which no HMA domain was identified by sequence analysis or structural prediction; Wrong annotation, constructs based on misannotated genes, resulting in missing or truncated HMA domains, thereby compromising Y2H assay.

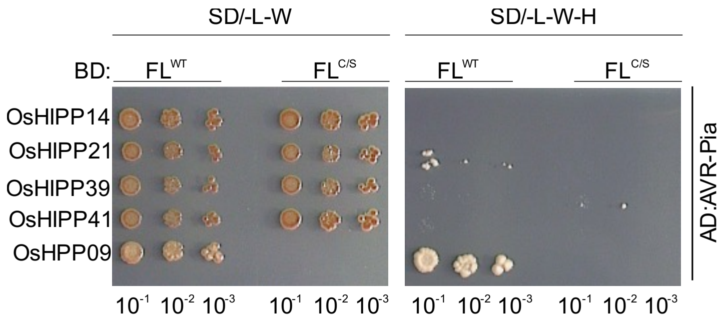


**Supplementary Fig. 2. Mutation of the isoprenylation motif of OsHIPPs does not promote AVR-Pia interaction.**

Y2H interaction analysis between AVR-Pia and either wildtype (WT) full-length (FL) OsHIPPs, including OsHIPP14, OsHIPP21, OsHIPP39 and OsHIPP41, or the corresponding FL OsHIPP isoprenylation mutants, in which the cysteine (C) residue was replaced by a non-prenylatable serine (S) residue. OsHPP09/AVR-Pia and was included as positive control. Serial dilutions of diploid yeast were spotted onto synthetic defined (SD) media to monitor growth (SD/-LW) or to assess protein-protein interactions (SD/-LWH). Photos were taken after 7 days of incubation. AD, activating domain; BD, binding domain. OsHIPP$^{C/S}$ mutants: OsHIPP14$^{C187S}$, OsHIPP21$^{C133S}$, OsHIPP39$^{C190S}$, OsHIPP41$^{C152S}$.

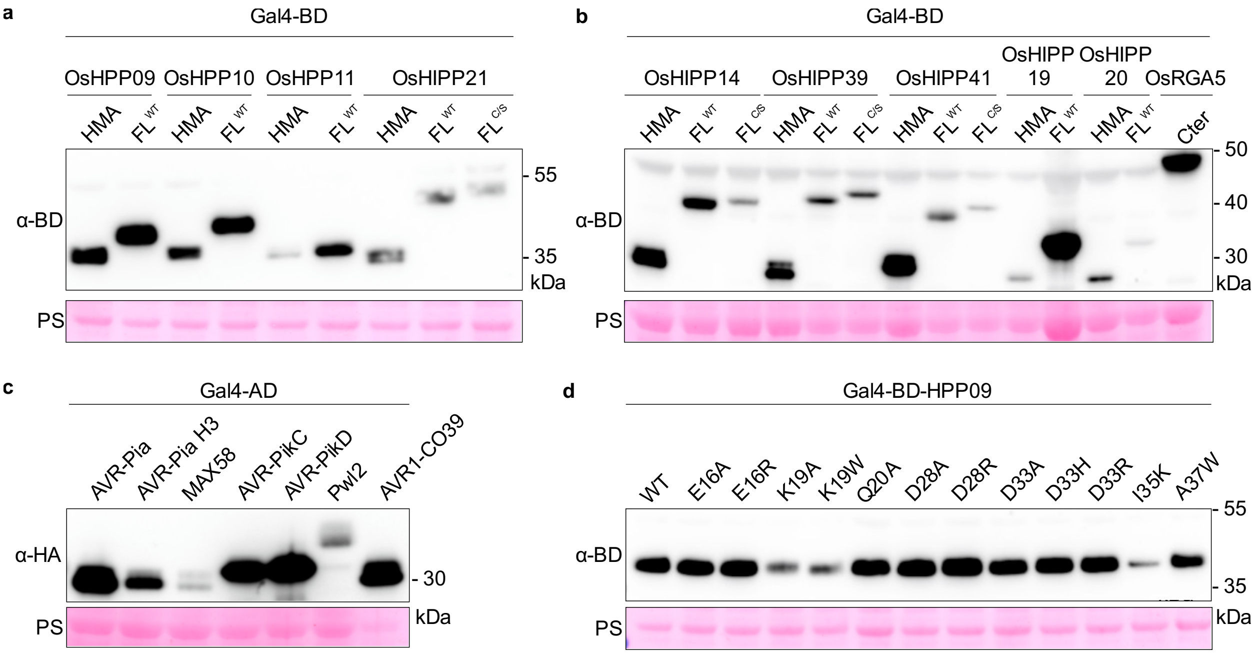


**Supplementary Fig. 3. Expression of bait and prey proteins in haploid yeast.**

**a** Total yeast protein extracts from haploid yeast were separated on a 10 – 20% Tricine SDS-PAGE gel and analysed by immunoblot using an anti-BD antibody to detect Gal4-BD fusion proteins with either the heavy metal-associated (HMA) domain, full-length (FL) OsH(I)PPs and the OsHIPP21 isoprenylation mutant (C/S). Protein loading is indicated by Ponceau S staining (PS). **b** Total yeast protein extracts from haploid yeast were separated on a 10% NuPAGE™ Bis-Tris gel and analysed by immunoblot using an anti-BD antibody to detect Gal4-BD fusion proteins with either the HMA domain, FL OsHIPPs and their corresponding isoprenylation mutants (C/S). Protein loading is indicated by Ponceau S staining (PS). **c** Total yeast protein extracts from haploid yeast were separated on a NuPAGE™ Bis-Tris gel and analysed by immunoblot using an anti-HA antibody to detect Gal4-AD fusion proteins with MAX effectors. Protein loading is indicated by Ponceau S staining (PS). **d** Total yeast protein extracts from haploid yeast were separated on a 10% NuPAGE™ Bis-Tris gel and analysed by immunoblot using an anti-BD antibody to detect Gal4-BD fusion proteins with FL OsHPP09 wildtype (WT) and mutant versions. Protein loading is indicated by Ponceau S staining (PS). OsHIPP$^{C/S}$ mutants: OsHIPP14$^{C187S}$, OsHIPP21$^{C133S}$, OsHIPP39$^{C190S}$, OsHIPP41$^{C152S}$.

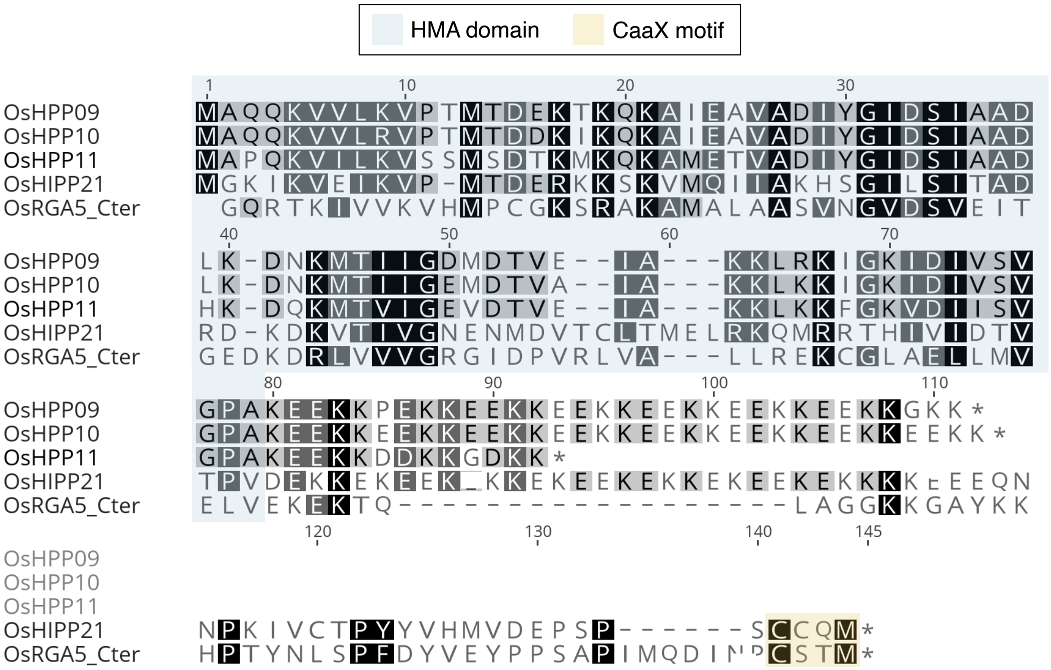


**Supplementary Fig. 4. Amino acid alignment of OsHPP09, OsHPP10, OsHPP11, OsHIPP21 and the C-terminus of OsRGA5.**

Amino acid alignment of full-length OsHPP09, OsHPP10, OsHPP11 and OsHIPP21 and residues 995-1116 of OsRGA5, including the HMA domain and the C-terminal region of the protein which concludes with a putative CaaX isoprenylation motif (CSTM). Sequence alignment was carried out with Clustal Omega. The blue shaded box highlights the HMA domain, while the yellow shaded box highlights the CaaX motif present in OsHIPP21 and OsRGA5.

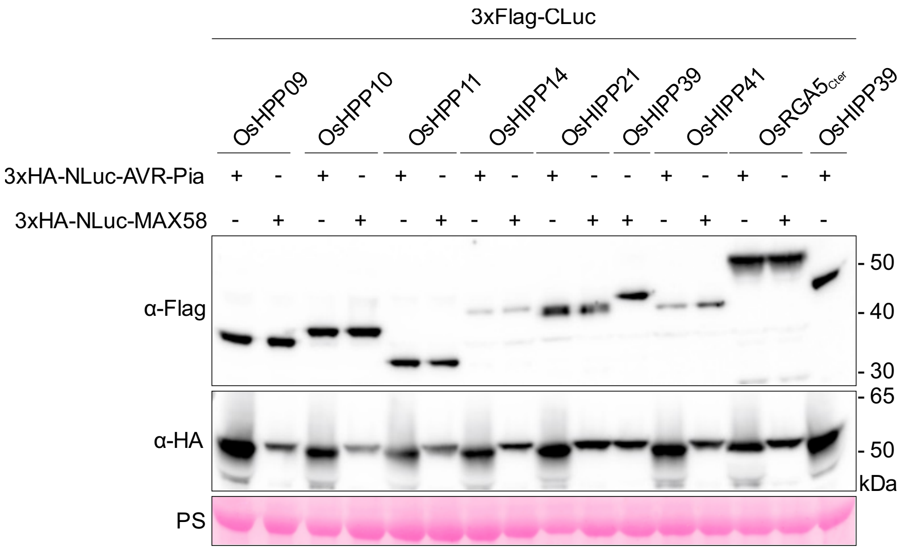


**Supplementary Fig. 5. Expression of CLuc- and NLuc fusion proteins (related to Fig. 1).**

Immunoblot analysis of transiently expressed proteins in *N. benthamiana*, including OsH(I)PPs N-terminally tagged with a 3xFlag epitope fused to the C-terminal part of luciferase (CLuc), and AVR-Pia and MAX58 N-terminally tagged with 3xHA and the N-terminal part of luciferase (NLuc). This experiment corresponds to replicate 1 of Fig. 1. Detection was performed using anti-Flag and anti-HA antibodies. Membrane was stripped after anti-Flag detection and re-probed with anti-HA. Protein loading is indicated by the Rubisco band visualized by Ponceau S. staining (PS).

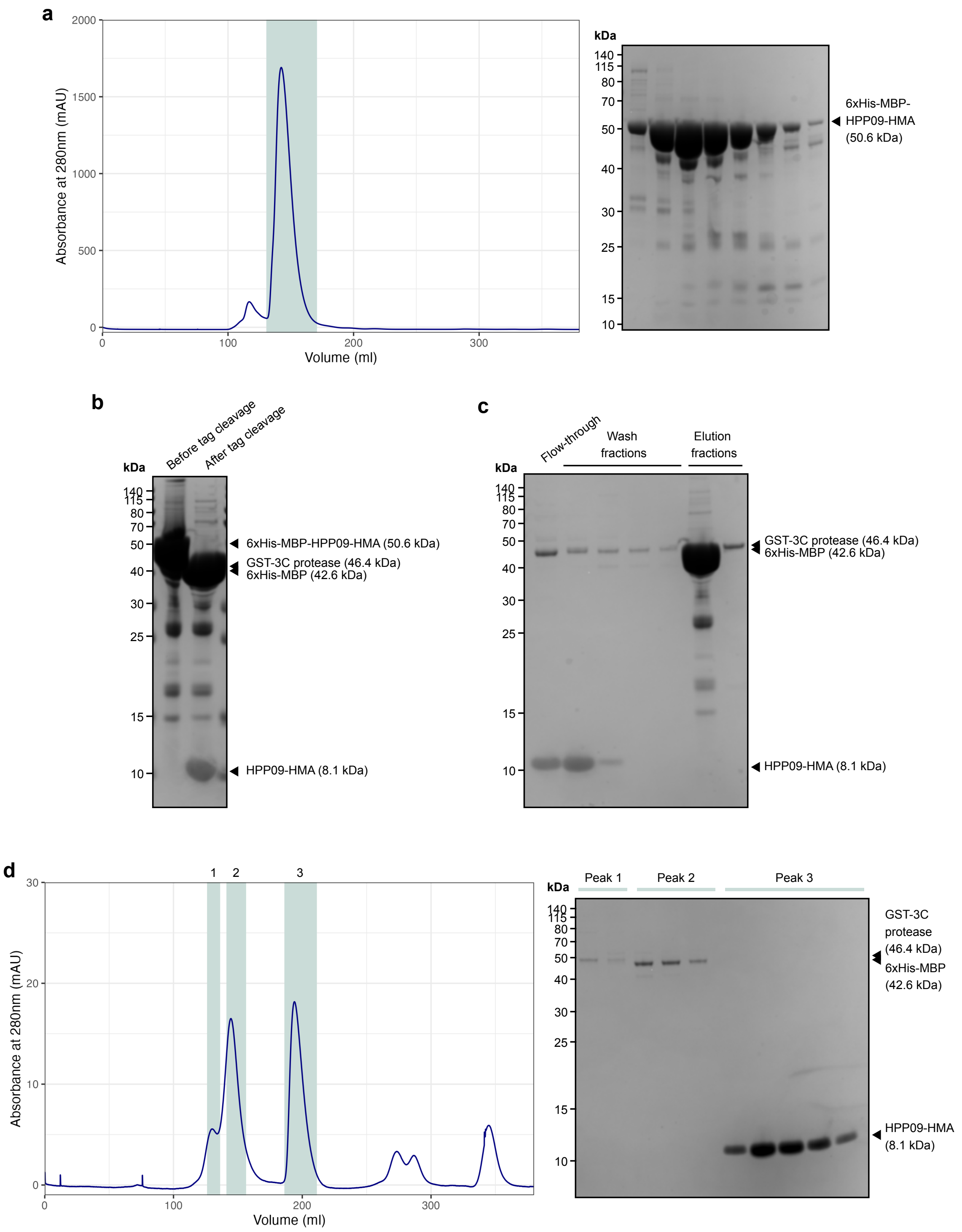


**Supplementary Fig. 6. Production and purification of OsHPP09-HMA.**

**a** Size exclusion chromatography elution trace for 6xHis-MBP-OsHPP09-HMA (following the initial IMAC purification step). The SDS-PAGE gel shows fractions corresponding to the peak in the shaded area. **b** SDS-PAGE gel of pooled fractions following size exclusion chromatography before and after cleavage of the 6xHis-MBP tag with GST-tagged 3C protease. **c** SDS-PAGE gel showing flow-through, wash and elution fractions from tandem HisTrapTM, MBPTrapTM and GSTrapTM affinity columns. **d** Size exclusion chromatography elution trace for OsHPP09-HMA. OsHPP09-HMA absorbs light at 280nm poorly (molar extinction coefficient of 1490 $M^{-1}cm^{-1}$). The SDS-PAGE gel shows fractions corresponding to the three peaks in the shaded areas. The peak elution volume of OsHPP09-HMA (193.9 ml) is consistent with dimerisation of the HMA domain.

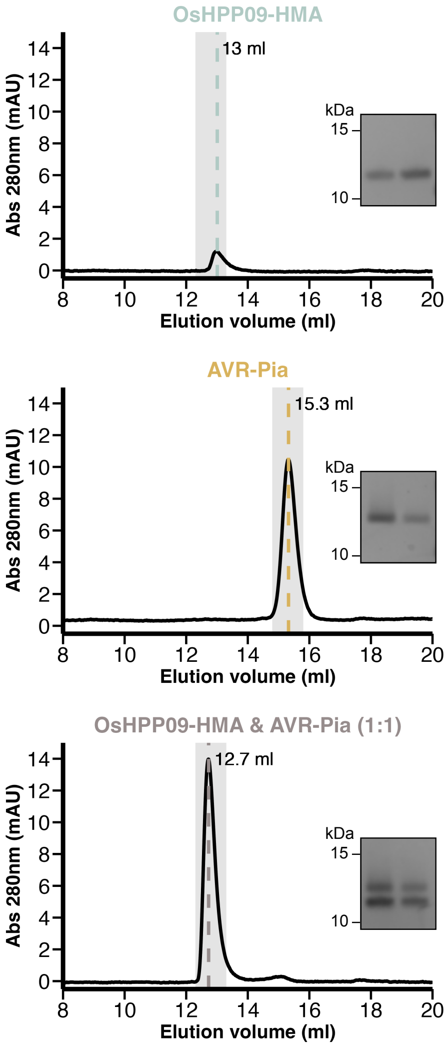


**Supplementary Fig. 7. AVR-Pia binds to the HMA domain of OsHPP09 in analytical size exclusion chromatography experiments.**

Analytical gel filtration traces obtained from injection of OsHPP09-HMA alone (top panel), AVR-Pia alone (middle panel) and the two proteins in a 1:1 molar ratio (bottom panel). Significant peaks are indicated by dashed lines with the elution volume labelled. SDS-PAGE gel inserts show fractions from the peak elution volumes indicated by the grey shaded regions. OsHPP09-HMA absorbs light at 280 nm poorly (molar extinction coefficient of 1490 M-1cm-1) so the peak corresponding to OsHPP09-HMA is small. The elution volume observed for OsHPP09-HMA is consistent with dimerisation, as has been observed for other purified HMA domains[2,3].

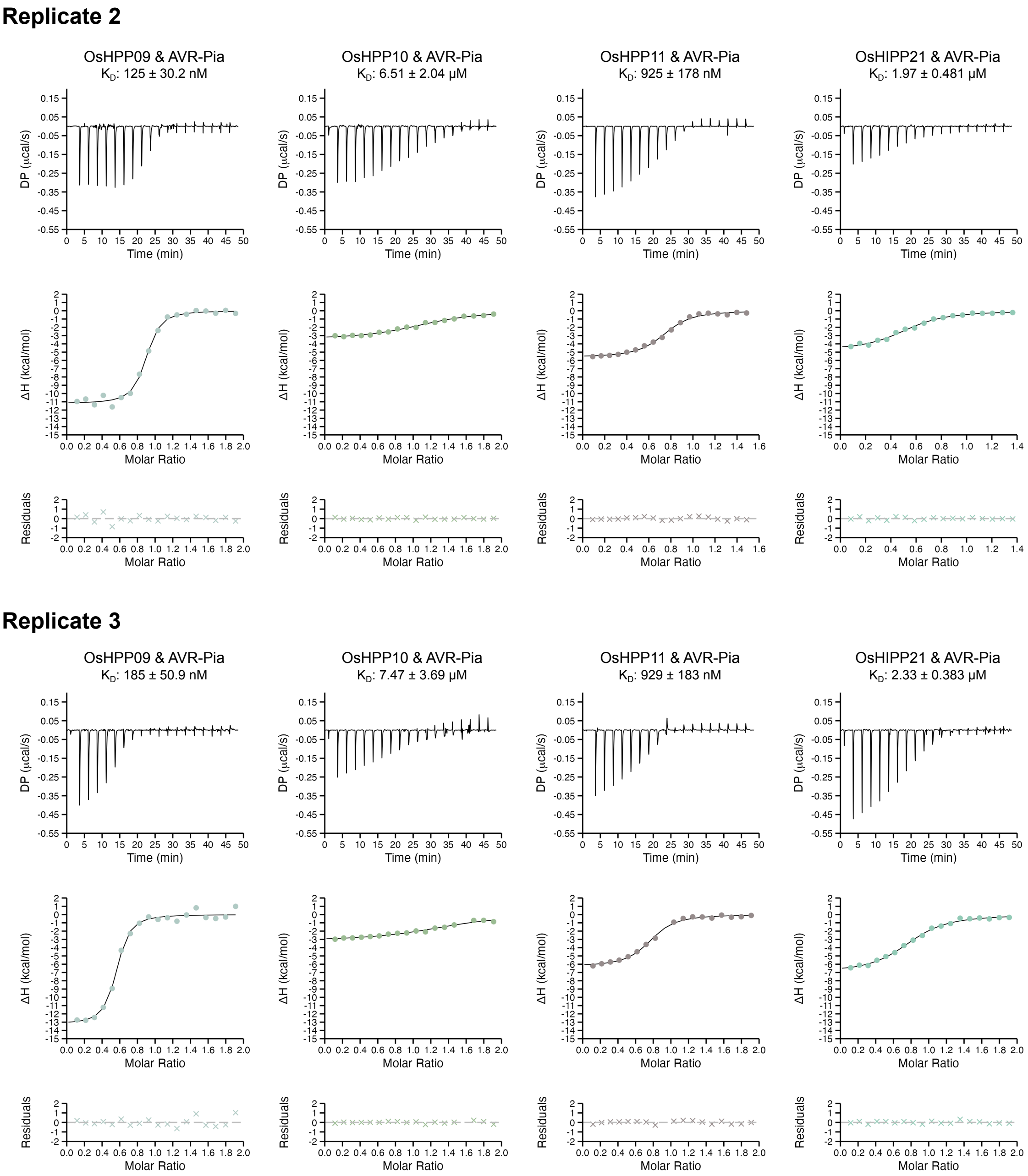


**Supplementary Fig. 8. Replicates of the ITC experiments presented in Fig. 2.**

Top panels show the raw thermograms obtained from titration of AVR-Pia into a solution containing the purified HMA domains. Central panels show the integrated heats (coloured dots) and binding isotherms fitted to a single site model (black line) using the MicroCal PEAQ-ITC analysis software (Malvern Panalytical). Bottom panels show the differences (coloured crosses) between the modelled and observed values (residuals).

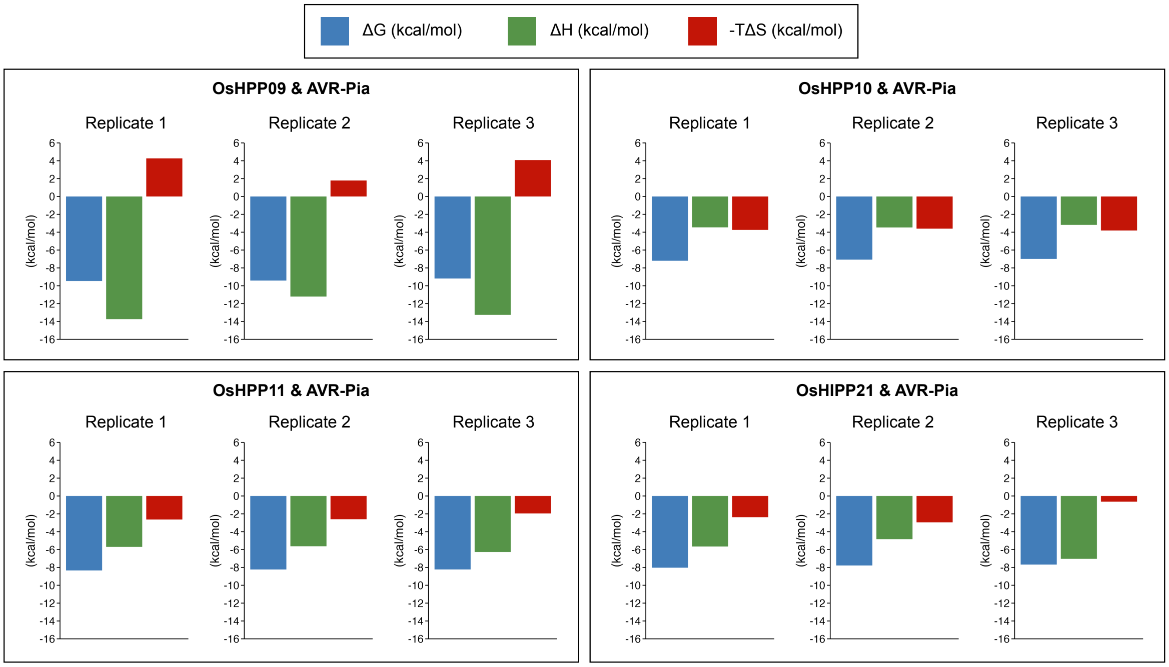


**Supplementary Fig. 9. Thermodynamic profiles from ITC experiments presented in Fig. 2 and Supplementary Fig. S8.**

Bars represent the magnitude of the determined thermodynamic parameters ΔG (blue bar), ΔH (green bar) and -TΔS (red bar).

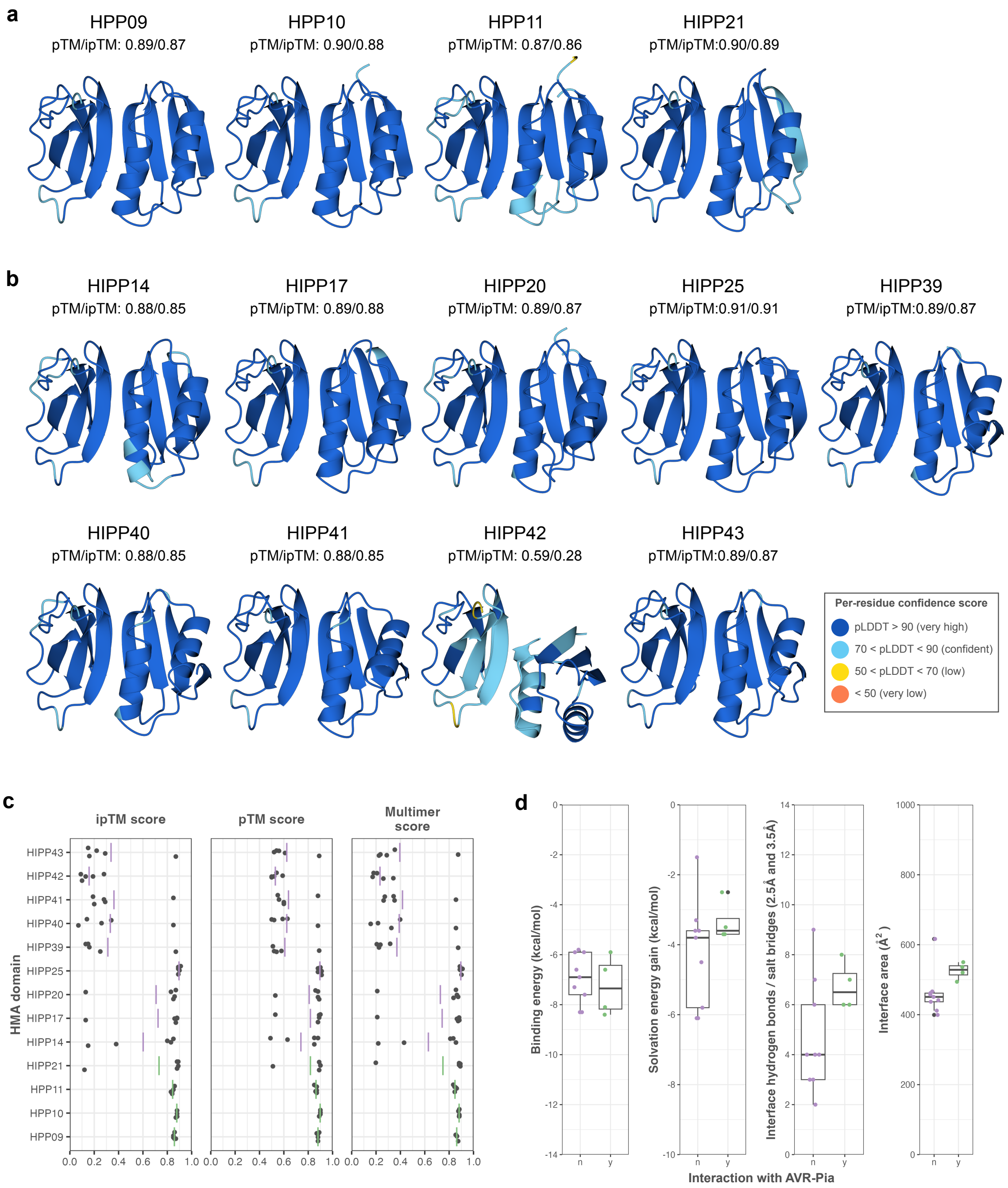


**Supplementary Fig. 10. AlphaFold2 models do not differentiate between HMA domains which interact with AVR-Pia and those which do not.**

AlphaFold2 (ColabFold v1.5.5) models of AVR-Pia in complex with HMA domains which **a** were experimentally shown to interact with AVR-Pia and **b** do not interact with AVR-Pia. Structure models are represented as ribbons and coloured by pLDDT score using the classical AlphaFold colour scheme as shown in the key. **c** Plots of ipTM, pTM and Multimer (0.8*iPTM + 0.2*pTM) scores for each of the five models generated for each HMA/AVR-Pia complex. Points indicate scores of individual models; green and purple lines indicate mean values for AVR-Pia-interacting and non-interacting HMA domains, respectively. **d** Comparison of interface parameters determined by qtPISA[4]) for the top ranked model of each HMA/AVR-Pia complex. Green and purple points indicate values for individual HMA/AVR-Pia models. The centre line of the box represents the median and the limits of the box represent the upper and lower quartiles. Whiskers extend to the smallest value within (Q1 − 1.5 × the interquartile range (IQR)) and the largest value within (Q3 + 1.5 × IQR).

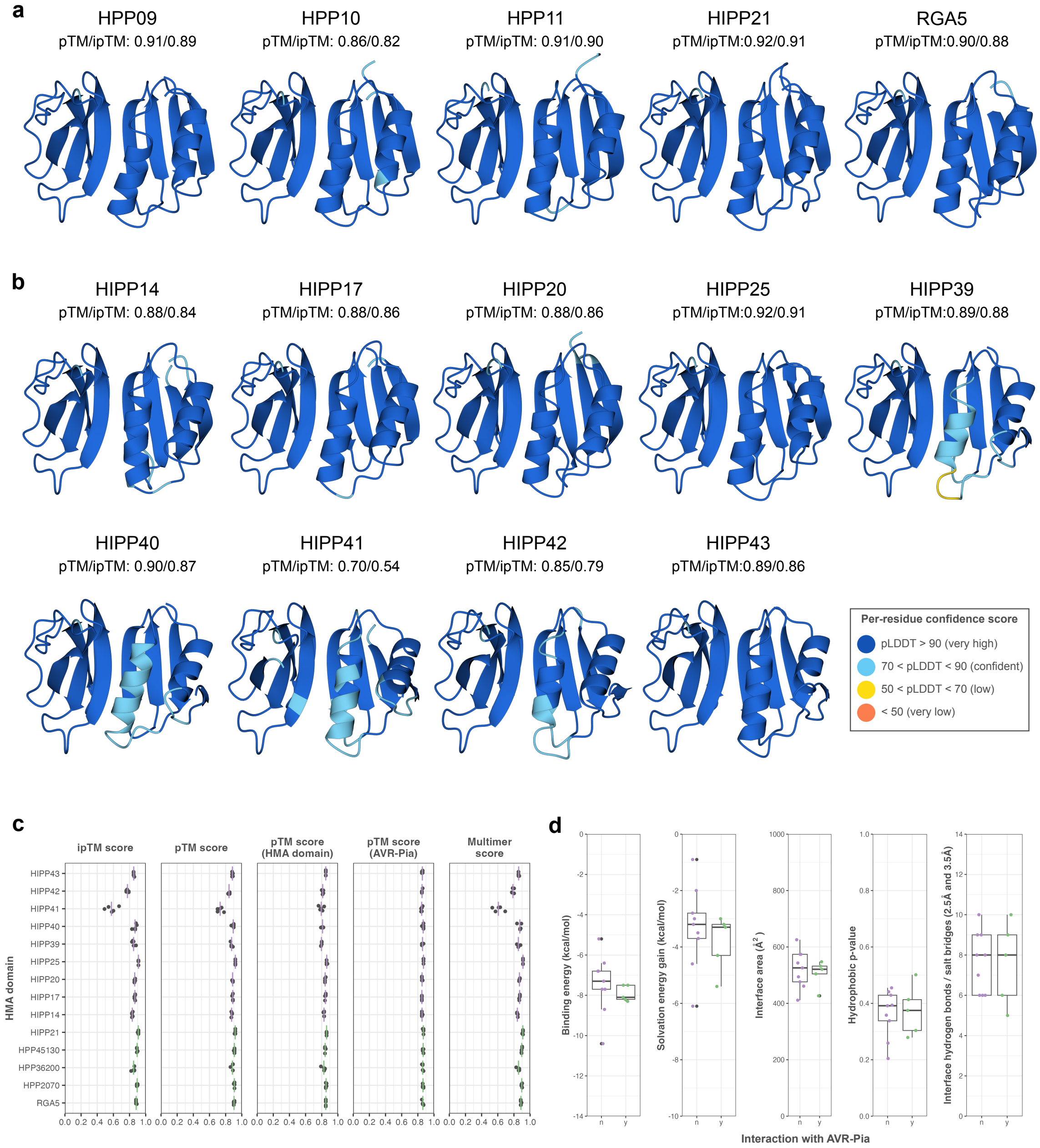


**Supplementary Fig. 11. AlphaFold3 models do not clearly differentiate between HMA domains which interact with AVR-Pia and those which do not.**

AlphaFold3 models of AVR-Pia in complex with HMA domains which **a** were experimentally shown to interact with AVR-Pia and **b** do not interact with AVR-Pia. Structure models are represented as ribbons and coloured by pLDDT score using the classical AlphaFold colour scheme as shown in the key. **c** Plots of ipTM, pTM and Multimer (0.8*iPTM + 0.2*pTM) scores for each of the top ranked models from each of the five seeds used to model each HMA/AVR-Pia complex. Points indicate scores of individual models; green and purple lines indicate mean values for AVR-Pia-interacting and non-interacting HMA domains, respectively. **d** Comparison of interface parameters determined by qtPISA[4]) for the top ranked model (selected from models from all five seeds) of each HMA/AVR-Pia complex. Green and purple points indicate values for individual HMA/AVR-Pia models. The centre line of the box represents the median and the limits of the box represent the upper and lower quartiles. Whiskers extend to the smallest value within (Q1 − 1.5 × the interquartile range (IQR)) and the largest value within (Q3 + 1.5 × IQR).

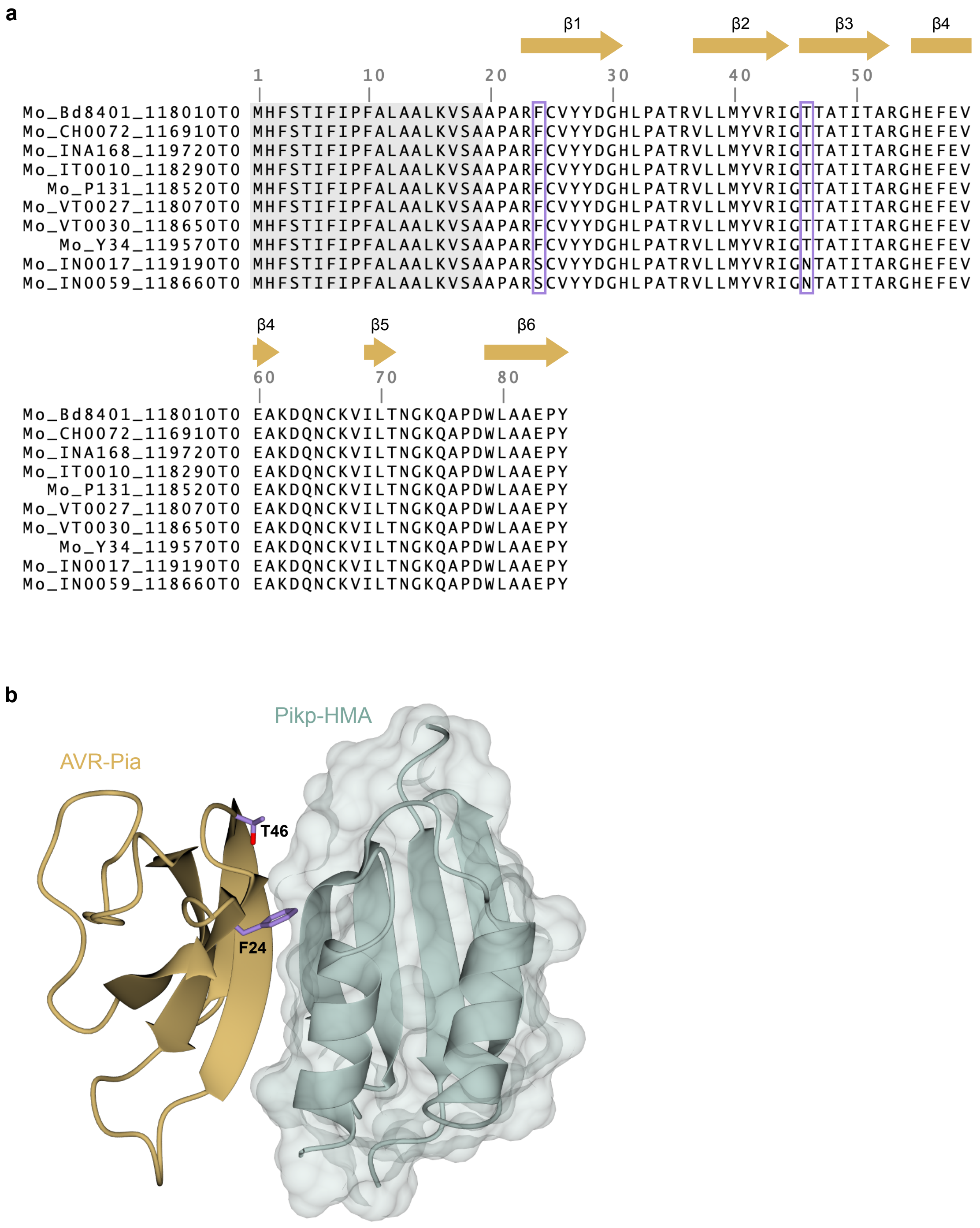


**Supplementary Fig. 12. AVR-Pia H3 differs from AVR-Pia in two amino acid positions.**

**a** Multiple sequence alignment of AVR-Pia variants from Magnaporthe oryzae isolates described in[5]. Signal peptide is highlighted by a shaded grey box. Polymorphic residues 24 (F, S) and 46

(T, N) are indicated in a purple outlined box. Arrows above the alignment indicate secondary structure elements. **b** The polymorphic residues F24 and T46 are located at the HMA binding interface. Crystal structure of AVR-Pia in complex with Pikp-HMA (PDB 6Q76[6]). AVR-Pia and Pikp-HMA are represented as gold and teal ribbons, respectively, with the side chains of AVR-Pia T46 and F24S shown in purple.

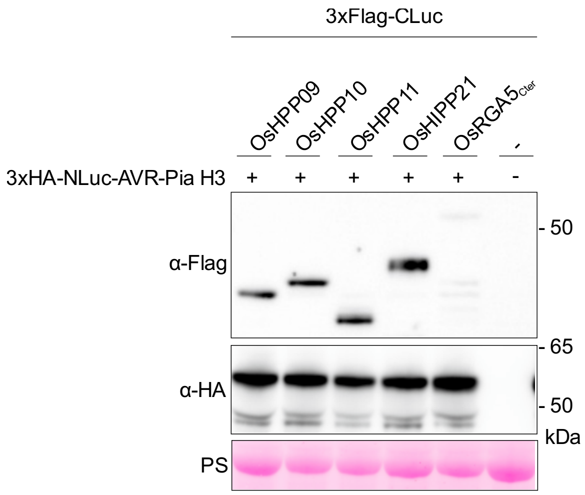


**Supplementary Fig. 13 Expression of CLuc- and NLuc fusion proteins (related to Fig. 3).**

Immunoblot analysis of transiently expressed proteins in *N. benthamiana*, including OsH(I)PPs and OsRGA5$_{Cter}$ (883 – 1116 aa) N-terminally tagged with a 3xFlag epitope fused to the C-terminal part of luciferase (CLuc), and AVR-Pia-H3 N-terminally tagged with 3xHA and the N-terminal part of luciferase (NLuc). This experiment corresponds to replicate 2 of **Fig. 3**. Detection was performed using anti-Flag and anti-HA antibodies. Membrane was stripped after anti-Flag detection and re-probed with anti-HA. Protein loading is indicated by the Rubisco band visualized by Ponceau S. staining (PS).

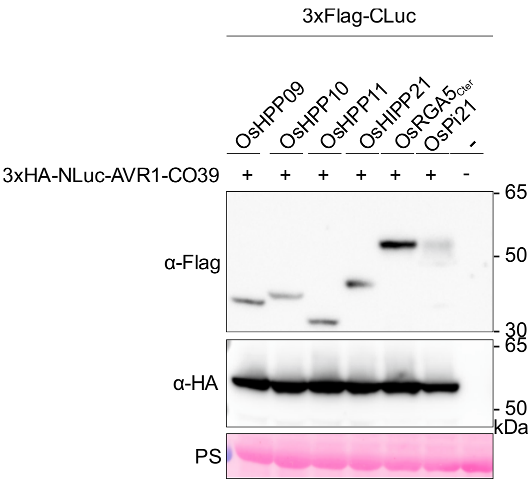


**Supplementary Fig. 14. Expression of CLuc- and NLuc fusion proteins (related to Fig. 4).**

Immunoblot analysis of transiently expressed proteins in *N. benthamiana*, including OsH(I)PPs and OsRGA5$_{Cter}$ (883 – 1116 aa) N-terminally tagged with a 3xFlag epitope fused to the C-terminal part of luciferase (CLuc), and AVR1-CO39 N-terminally tagged with 3xHA and the N-terminal part of luciferase (NLuc). This experiment corresponds to replicate 1 of **Fig. 4**. Detection was performed using anti-Flag and anti-HA antibodies. Membrane was stripped after anti-Flag detection and re-probed with anti-HA. Protein loading is indicated by the Rubisco band visualized by Ponceau S. staining (PS).

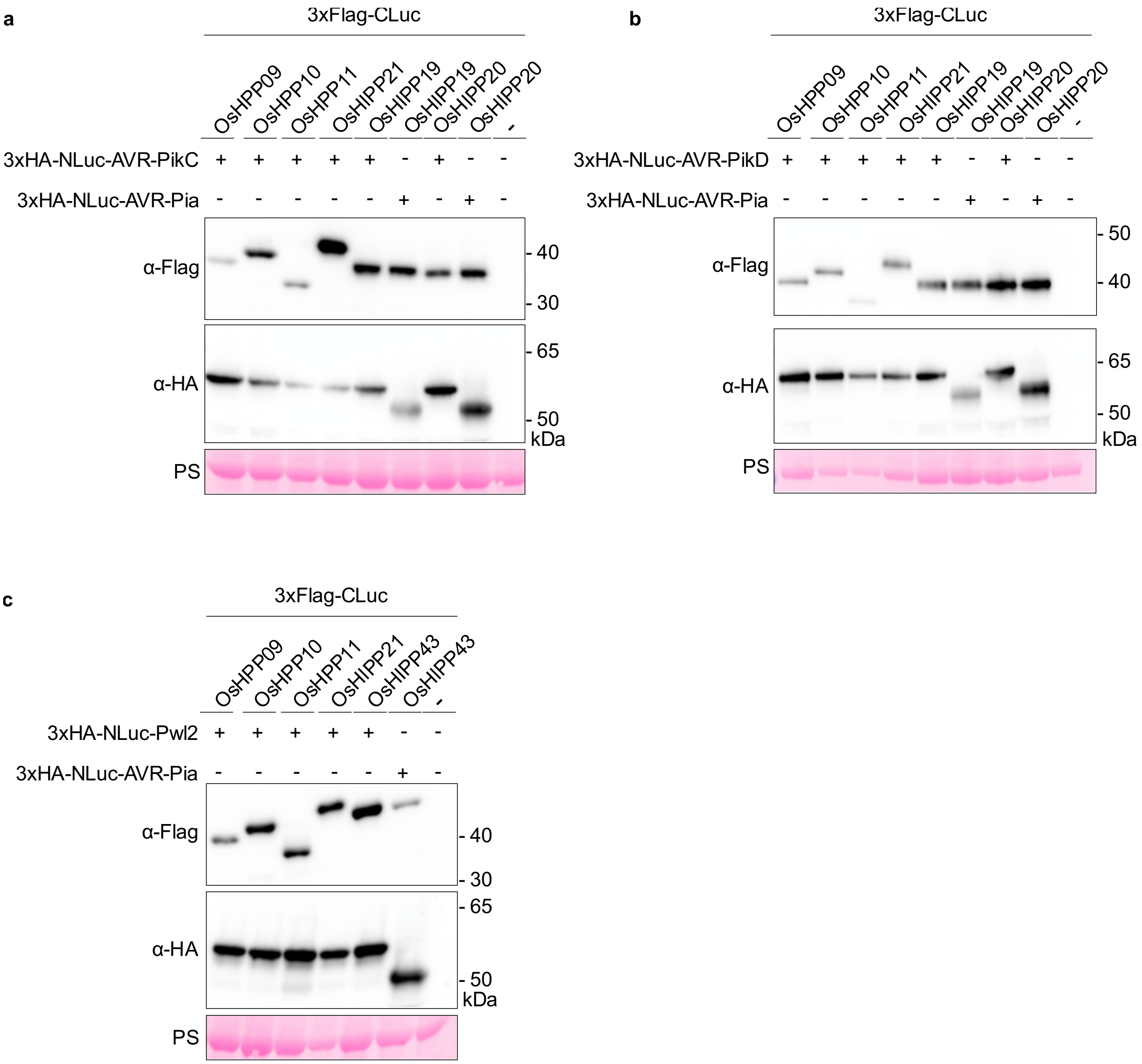


**Supplementary Fig. 15. Expression of CLuc- and NLuc fusion proteins (related to Fig. 5).**

Immunoblot analysis of transiently expressed proteins in *N. benthamiana*, including OsH(I)PPs N-terminally tagged with a 3xFlag epitope fused to the C-terminal part of luciferase (CLuc), and AVR-Pia, AVR-PikC, AVR-PikD and Pwl2 N-terminally tagged with 3xHA and the N-terminal part of luciferase (NLuc). This experiment corresponds to replicate 1 of **Fig. 5**. Detection was performed using anti-Flag and anti-HA antibodies. Membrane was stripped after anti-Flag detection and re-probed with anti-HA. Protein loading is indicated by the Rubisco band visualized by Ponceau S. staining (PS).

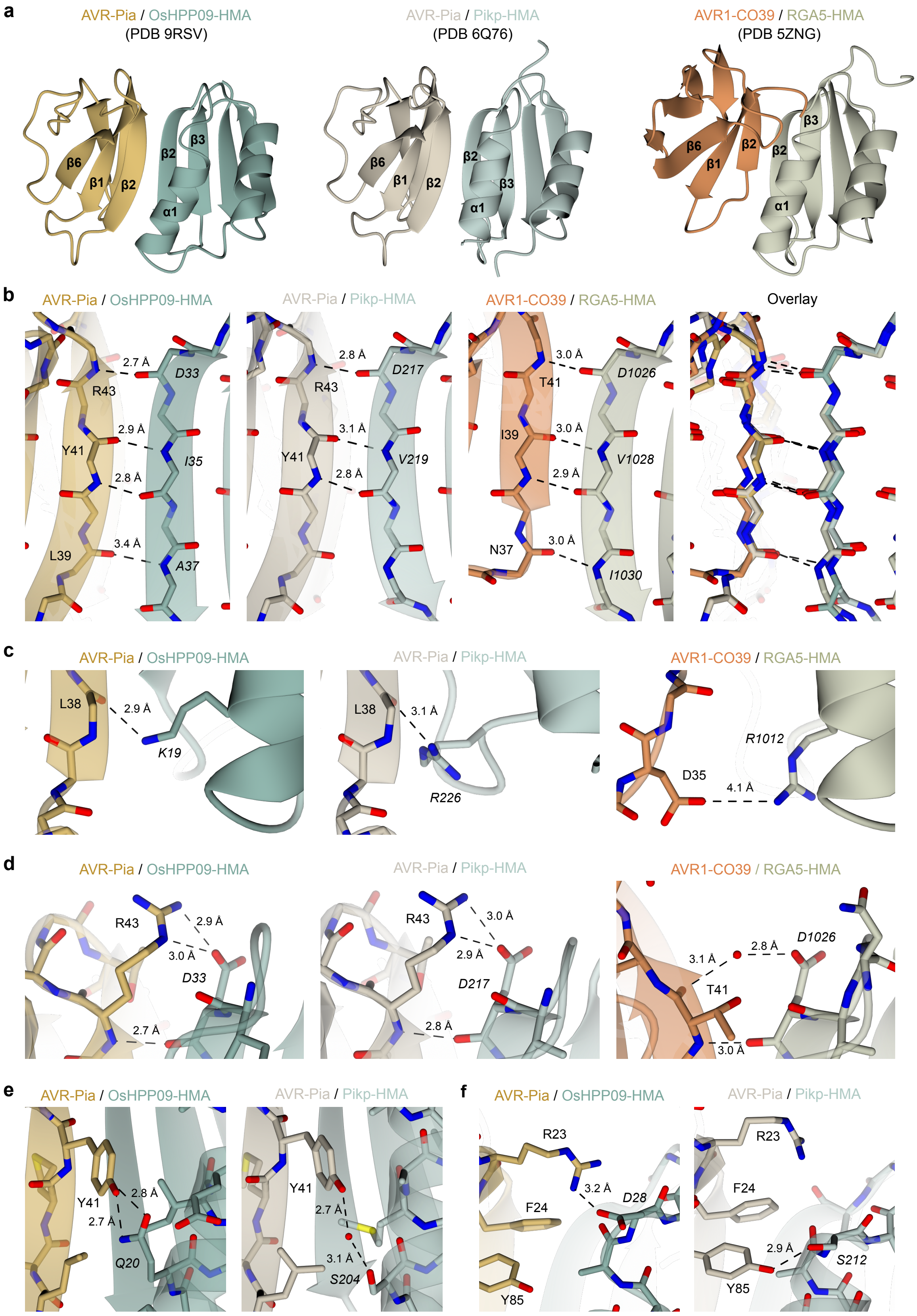
a
AVR-Pia / OsHPP09-HMA
(PDB 9RSV)
AVR-Pia / Pikp-HMA
(PDB 6Q76)
AVR1-CO39 / RGA5-HMA
(PDB 5ZNG)
β6
β1
β2
β3
α1
b
AVR-Pia / OsHPP09-HMA
AVR-Pia / Pikp-HMA
AVR1-CO39 / RGA5-HMA
Overlay
2.7 Å
D33
R43
2.9 Å
Y41
I35
2.8 Å
3.4 Å
L39
A37
2.8 Å
D217
3.1 Å
V219
3.0 Å
T41
D1026
I39
V1028
2.9 Å
N37
I1030
c
L38
K19
R226
R1012
D35
4.1 Å
d
3.0 Å
D1026
e
Q20
S204
f
R23
F24
Y85
3.2 Å
D28
S212
2.9 Å

**Supplementary Fig. 16. Comparison of binding interfaces in the crystal structures of AVR-Pia in complex with OsHPP09-HMA (PDB accession code 9RSV), AVR-Pia/Pikp-HMA (6Q76[6]) and AVR1-CO39/RGA5-HMA (5ZNG[2])**

**a** Global interface of the effector/HMA complex. Structures are represented as gold (AVR-Pia), teal (OsHPP09-HMA), brown (AVR-Pia), pale blue (Pikp-HMA), orange (AVR1-CO39) and green (RGA5-HMA) ribbons with relevant secondary structure elements indicated. **b** Hydrogen bonds between residues in β2 of the effector and β2 of the HMA domain. Main chain atoms are represented as cylinders (side chains not shown for clarity). Hydrogen bonds are represented as black dashed lines with lengths (determined by qtPISA) indicated. **c-f** Comparison of residues involved in forming intermolecular contacts at the effector/HMA interface. Structures are presented in ribbon representation with relevant residues shown as cylinders. Hydrogen bonds are represented as black dashed lines with lengths (determined by qtPISA) indicated. Water molecules are represented as red spheres.

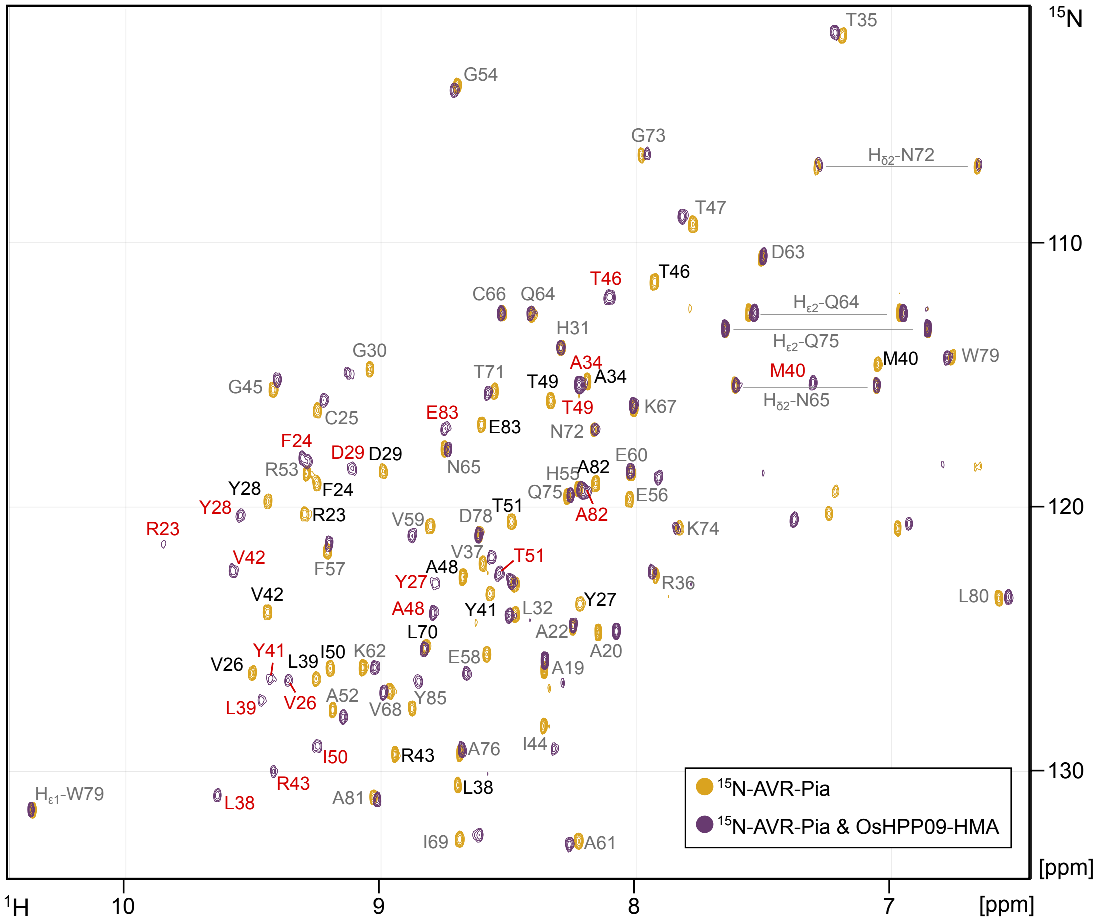


**Supplementary Fig. 17. Overlay of HSQC spectra of $^{15}$N-AVR-Pia (40 μM) in the absence (orange cross-peaks) and presence (purple cross-peaks) of unlabelled OsHPP09-HMA (40 μM).**

Grey cross-peak labels are used for residues with little or no chemical shift perturbation. For substantial chemical shifts, black cross-peak labels correspond to spectra for $^{15}$N-AVR-Pia alone (previously assigned[7]), and red cross-peak labels correspond to spectra for $^{15}$N-AVR-Pia in the presence of OsHPP09-HMA.

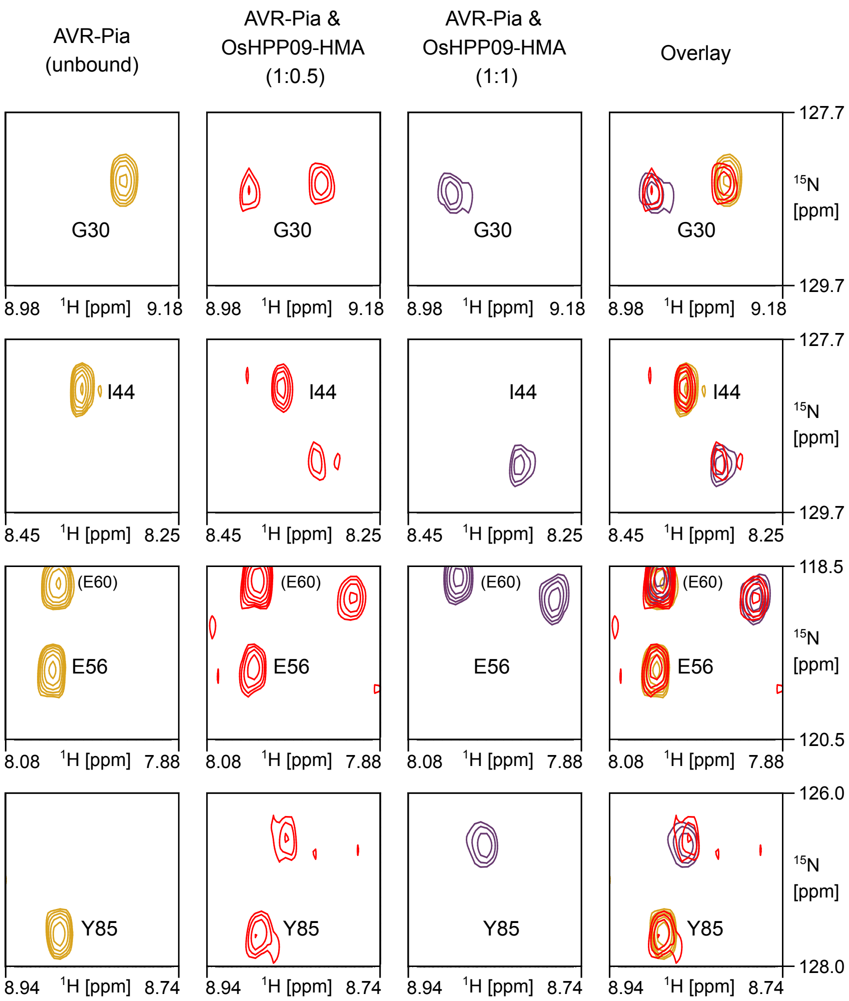


**Supplementary Fig. 18. The AVR-Pia/OsHPP09-HMA complex is in slow exchange.**

Cross-peaks corresponding to residues G30, I44, E56 and Y85 extracted from [$^1$H,$^{15}$N] HSQC spectra from NMR titrations performed with $^{15}$N-labelled AVR-Pia and OsHPP09-HMA at molar ratios of 1:0 (free AVR-Pia; orange cross-peaks), 1:0.5 (intermediate; red cross-peaks) and 1:1 (bound AVR-Pia; purple cross-peaks). The final column shows the overlay of the different spectra.

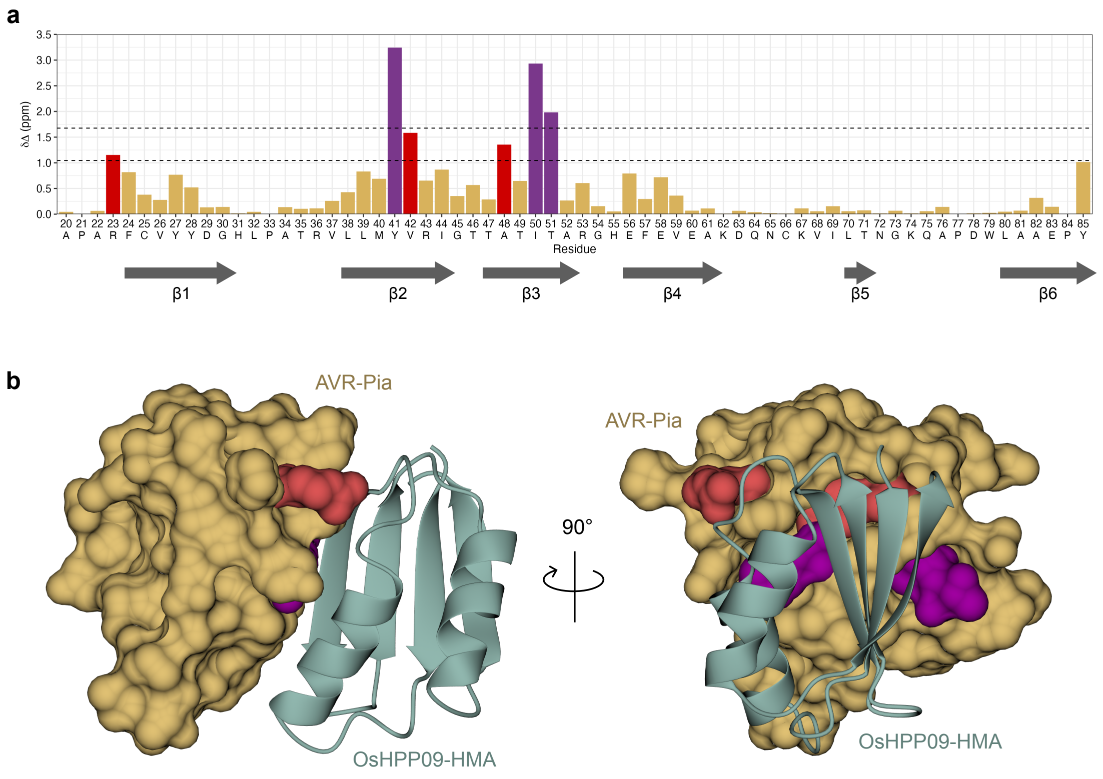


**Supplementary Fig. 19. The OsHPP09-HMA binding interface involves residues in β2 and β3 of AVR-Pia.**

**a** Barplot of $^{1}H/^{15}N$ chemical shift perturbations (Δδ) for the amide groups of AVR-Pia. Dashed lines represent Δδ thresholds of $\overline{\Delta\delta}$ + 1σ and $\overline{\Delta\delta}$ + 2σ. Significant chemical shift variations (> $\overline{\Delta\delta}$ + 1σ or > $\overline{\Delta\delta}$ + 2σ) are indicated in red and purple, respectively. **b** Residues giving significant chemical shift variations of > $\overline{\Delta\delta}$ + 1σ or > $\overline{\Delta\delta}$ + 2σ represented in red or purple, respectively, on the crystal structure (PDB 9RSV) of AVR-Pia (surface representation) bound to OsHPP09-HMA (teal ribbon representation). The two views are rotated by 90° as indicated.

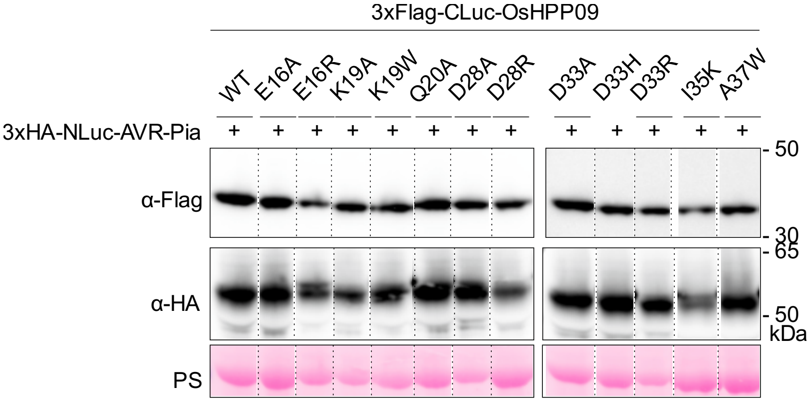


**Supplementary Fig. 20. Expression of CLuc- and NLuc fusion proteins (related to Fig. 7).**

Immunoblot analysis of transiently expressed proteins in *N. benthamiana*, including OsHPP09 wildtype (WT) and mutant versions N-terminally tagged with a 3xFlag epitope fused to the C-terminal part of luciferase (CLuc), and AVR-Pia N-terminally tagged with 3xHA and the N-terminal part of luciferase (NLuc). This experiment corresponds to replicate 2 of Fig. 7. Detection was performed using anti-Flag and anti-HA antibodies. Membrane was stripped after anti-Flag detection and re-probed with anti-HA. Protein loading is indicated by the Rubisco band visualized by Ponceau S. staining (PS). Dashed lines between bands indicate that lanes were cut from the same blot but were non-adjacent.

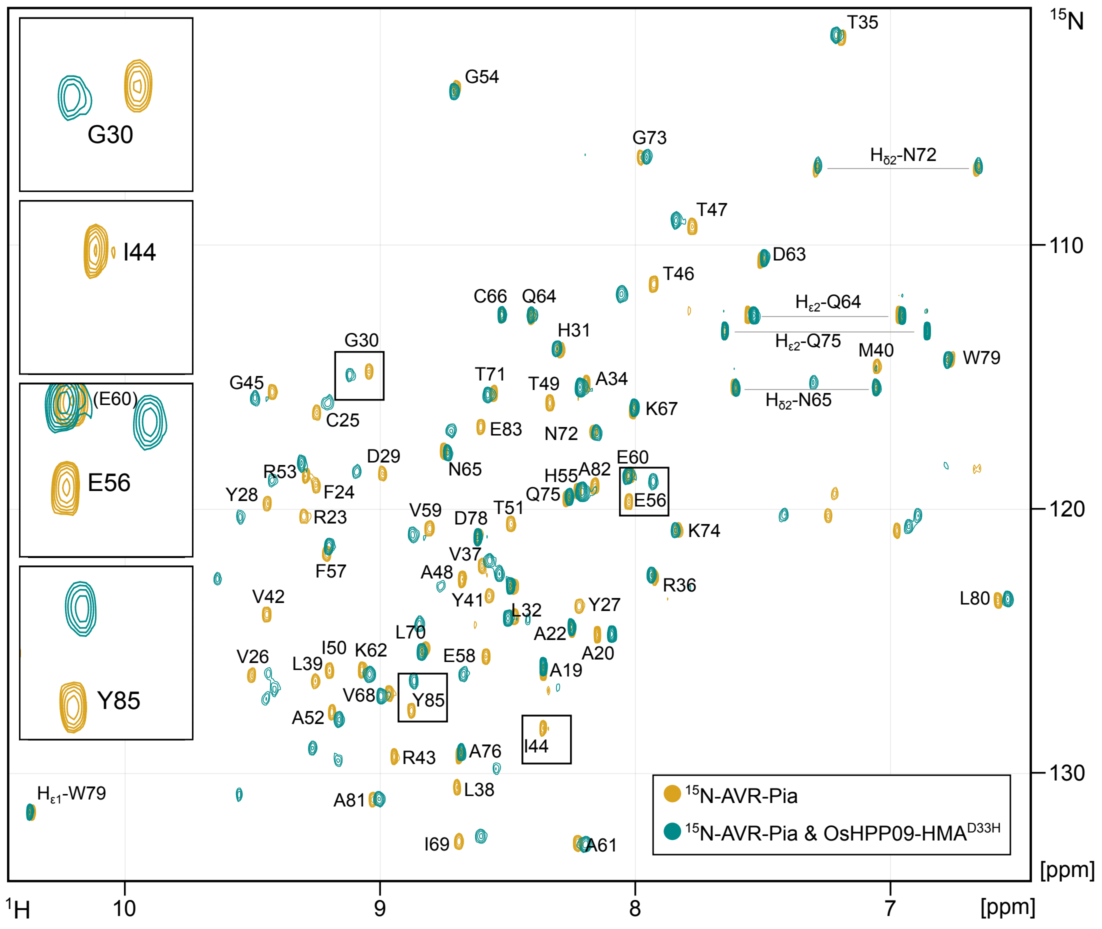

**Supplementary Fig. 21. HSQC spectra of $^{15}$N-AVR-Pia (40 µM) in the absence (orange cross-peaks) and presence (turquoise cross-peaks) of unlabelled OsHPP09$^{D33H}$-HMA (40 µM).**

Cross-peak labels correspond to the $^{15}$N-AVR-Pia spectra (previously assigned[7]). Zoom inset panels show cross-peaks corresponding to residues G30, I44, E56 and Y85 to illustrate the chemical shift perturbations between the free ($^{15}$N-AVR-Pia alone) and bound ($^{15}$N-AVR-Pia with OsHPP09$^{D33H}$-HMA) as presented in Fig. 7. Zoomed areas are indicated by boxes on the spectra.

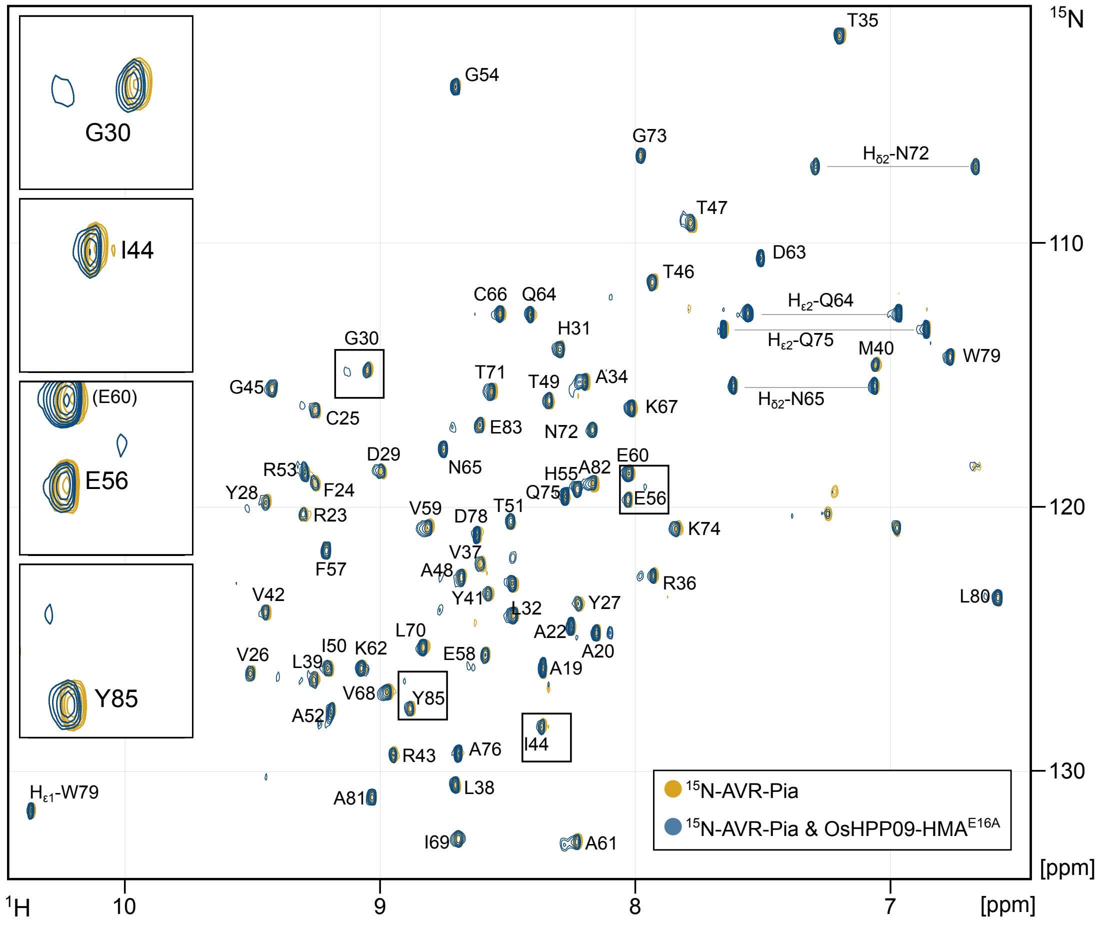

**Supplementary Fig. 22. HSQC spectra of $^{15}$N-AVR-Pia (40 µM) in the absence (orange cross-peaks) and presence (blue cross-peaks) of unlabelled OsHPP09$^{E16A}$-HMA (40 µM).**

Cross-peak labels correspond to the $^{15}$N-AVR-Pia spectra (previously assigned[7]). Zoom inset panels show cross-peaks corresponding to residues G30, I44, E56 and Y85 to illustrate the chemical shift perturbations between the free ($^{15}$N-AVR-Pia alone) and bound ($^{15}$N-AVR-Pia with OsHPP09$^{E16A}$-HMA) as presented in Fig. 7. Zoomed areas are indicated by boxes on the spectra.

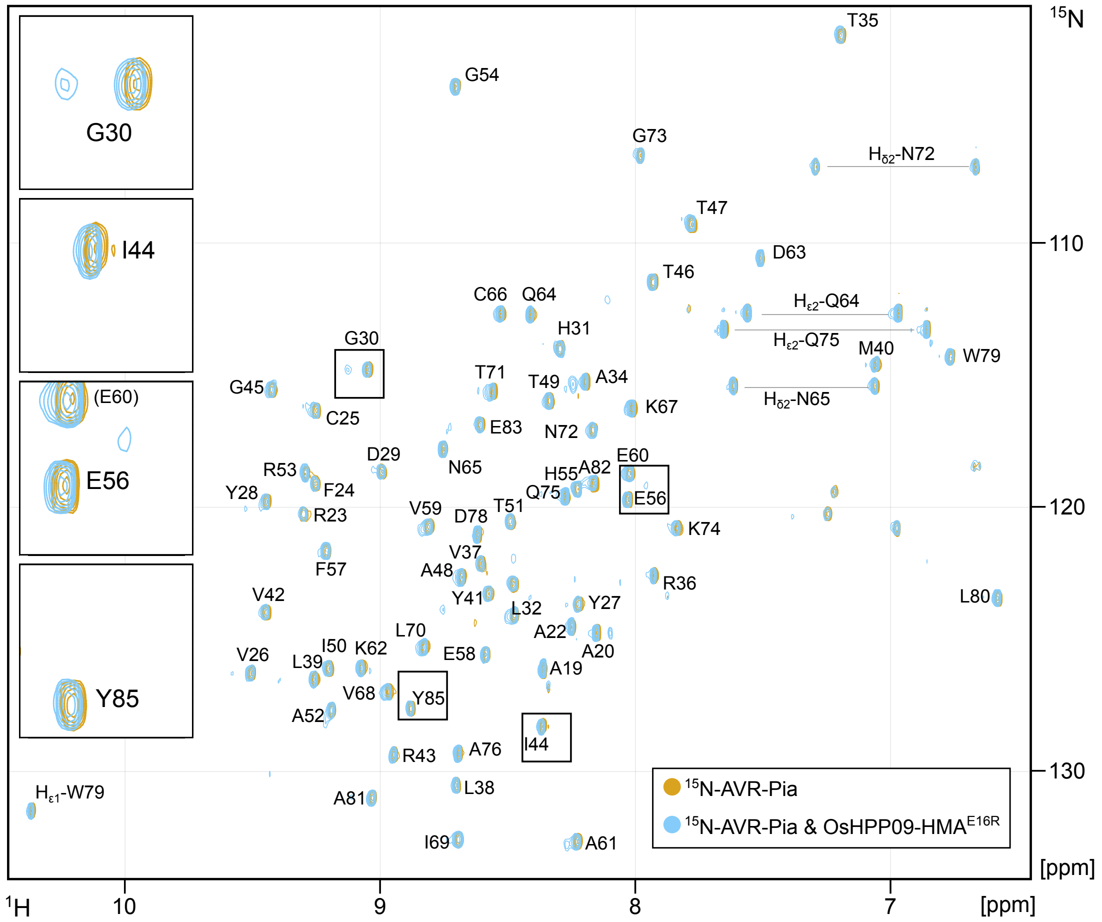

**Supplementary Fig. 23. HSQC spectra of $^{15}$N-AVR-Pia (40 µM) in the absence (orange cross-peaks) and presence (pale blue cross-peaks) of unlabelled OsHPP09$^{E16R}$-HMA (40 µM).**

Cross-peak labels correspond to the $^{15}$N-AVR-Pia spectra (previously assigned[7]). Zoom inset panels show cross-peaks corresponding to residues G30, I44, E56 and Y85 to illustrate the chemical shift perturbations between the free ($^{15}$N-AVR-Pia alone) and bound ($^{15}$N-AVR-Pia with OsHPP09E16R-HMA) as presented in Fig. 7. Zoomed areas are indicated by boxes on the spectra.

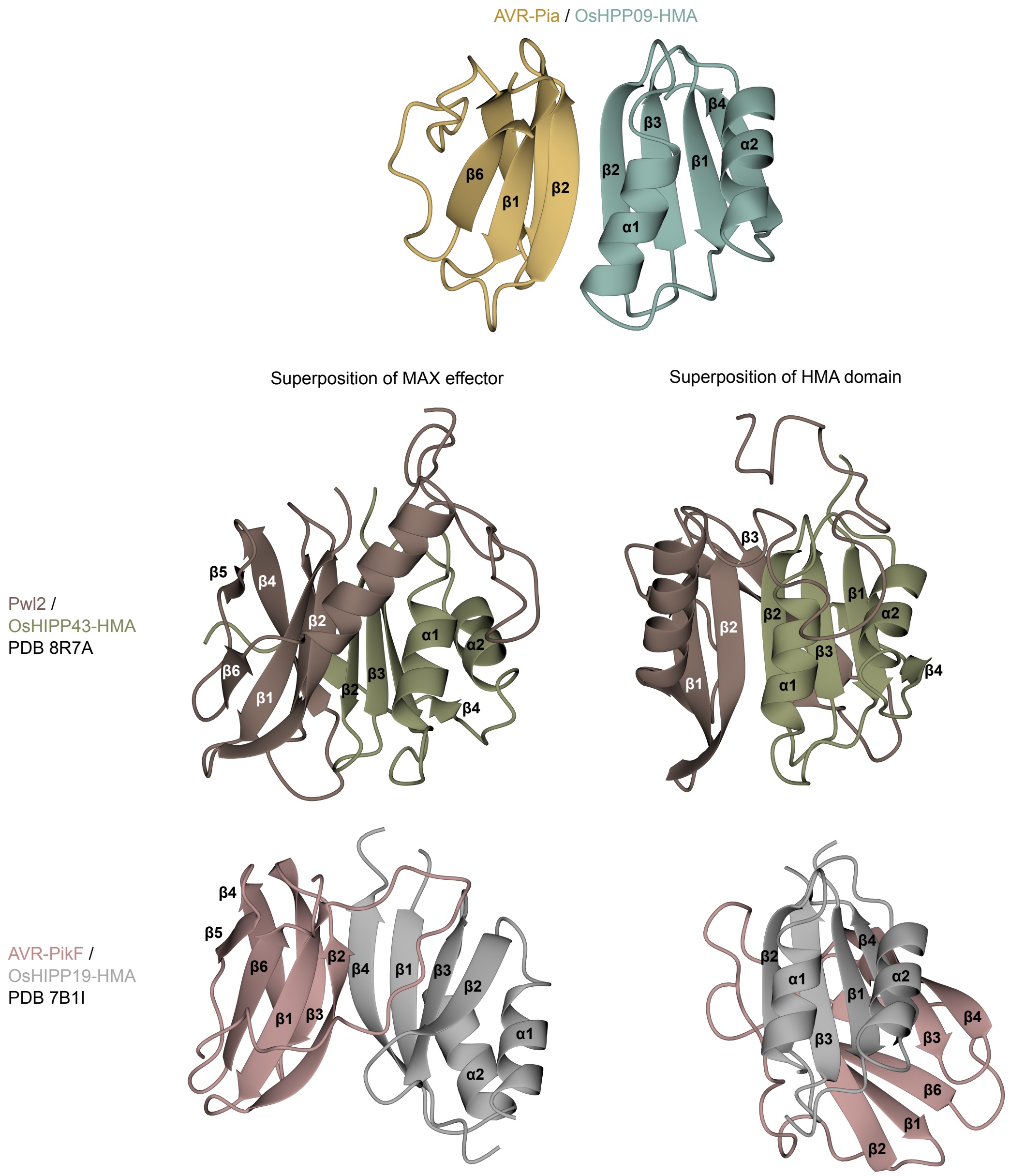


**Supplementary Fig. 24. Comparison of binding interfaces in the crystal structures of MAX effectors bound to HMA domains of H(I)PPs.**

The structures of AVR-Pia in complex with OsHPP09-HMA (PDB accession code 9RSV), AVR-PikF in complex with OsHIPP19-HMA (PDB accession code 7B1I[8]) and Pwl2 in complex with OsHIPP43-HMA (PDB accession code 8R7A[9]) are represented as gold (AVR-Pia), teal (OsHPP09-HMA), brown (Pwl2), green (OsHIPP43-HMA), pink (AVR-PikF) and grey (OsHIPP19-HMA) ribbons with relevant secondary structure elements indicated. Superposition of MAX effectors (left) and HMA domains (right) was carried out in CCP4mg using secondary structure matching.

**Supplementary Table S1. Amino acid sequences and MSU/RGAP and RAP-DB/IRGSP identifiers for the H(I)PPs used in the present study.**

| H(I)PP name | MSU/RGAP[1] ID | RAP-DB/IRGSP[2] ID | Amino acid sequence[3] |
|---|---|---|---|
| OsHPP09 | LOC_Os03g02070.1 | Os03t0111400-01 | **MAQQKVVLKVPTMTDEKTKQKAIEAVADIYGIDSIAADLKDNKMTIIGDMDTVEIAKKLRKIGKIDIVSVGPA**KEEKKPEKKEEKKEEKKEEKKEEKKEEKKGKK* |
| OsHPP10 | LOC_Os10g36200.2 | [Os10t0506100-01][4] | **MAQQKVVLRVPTMTDDKIKQKAIEAVADIYGIDSIAADLKDNKMTIIGEMDTVAIAKKLKKIGKIDIVSVGPA**KEEKKEEKKEEKKEEKKEEKKEEKKEEKKEEKK* |
| OsHPP11 | LOC_Os04g45130.1 | [Os04t0533900-01][5] | **MAPQKVILKVSSMSDTKMKQKAMETVADIYGIDSIAADHKDQKMTVIGEVDTVEIAKKLKKFGKVDIISVGPA**KEEKKDDKKGDKK* |
| OsHIPP05 (Pi21) | LOC_Os04g32850.1 | Os04t0401000-01 | **MGILVILVDLQCCRCDAKIRKVLGCLEEEYCIEKVEYDVKNNRVIVRGKFDPEKLCKKIWCKAGKIIKEILIVD**VWPPPLPQPPPPCKPPPCEKPPEDCKPKPCHCCSCEKPKPKPKPCHCEKPKPCHCEKPKPCEKPPPCKPEEPPKPPPEKPPPKPECKLVPYPYPVPYPYAGQWCCPKPEPPKPPPEPPKEPEPPKPCGCSHAFVCVCKPAPPPPPPCGCSGGHGNCGCGIRPWPPQVWPPPPVCPPPPWCYTEDNANACSIM* |
| OsHIPP21 | LOC_Os09g09830.1 | Os09t0271100-02 | **MGKIKVEIKVPMTDERKKSKVMQIIAKHSGILSITADRDKDKVTIVGNENMDVTCLTMELRKQMRRTHIVIDTVTPV**DEKKEKEEKEKKEKEEKEKKEKEEKEKKKKEEEQNNPKIVCTPYYVHMVDEPSPSCCQM* |
| OsHIPP14 | LOC_Os04g39320.4 | Os04t0468600-02 | MR**KEIVIRLQSSEKGHKKAIKVAAAVSGVESVTLAGEDKNLLLVIGFGVDSNDLTEKLRRKVGHAEVVELRTV**DADELMRVAAANQYPYRYYPGAPPPAPYYGNGGYPPPHQRGGGGGGSGGGYYTPMTMATGGYYGGGGGGYPQYGQSSSYPQYGQSSSYYPPAAAATTNTHTVVHHQYANNDPDSCAIM* |
| OsHIPP19 | LOC_Os04g39350.1 | Os04t0469000-01 | **MKQKIVIKVSMPCEKSRSKAMKLVVMASGVSSVEVTGDGKDRLQVVGDGVDAACLVTCLRKKIGHAELVQVEEVKEK**KPEEKKPEEKKPEPCYCPHPCYYHHHYGGIPVAVGDQPSDPCSIM* |
| OsHIPP39 | LOC_Os03g02860.1 | Os03t0120400-01 | MSTVSSALSSFLYCCFSPTGGHRHGHRAGAYYYSSHPTSTNTYYYEGGLAGRRMGRSRPLS**LQTVELKVRMCCSGCERVVKHALMKLRGVDSVEVELEMEKVTVTGYVERQRVLKEVRRAGKKAEFW**PNPDLPLYFTSAKDYFHDEESFRPSYNYYRHGYNGDKHGHLPEPHRGADPVSNLFNDDDVNACSIM* |
| OsHIPP41 | LOC_Os03g06080.1 | Os03t0156600-01 | MGVDDIIAELRVLPAKILLKKKPKQF**QKVEVKVRMDCEGCERKVRKAVEEMKGVSSVEVDAKQNKVTVTGYVEQEEVVGRLRRRAGKKAEPW**PYVPYDVVPHPYAPGAYDKKAPPGYVRNALADPDAAPLARATEEEEKLASAFSDENPNSCAVM* |
| OsHIPP43 | LOC_Os01g32330.1 | Os01t0507700-01 | MGVLDSLSDMCSLTETKEALKLRKKRP**LQTVNIKVKMDCEGCERRVKNAVKSMRGVTSVAVNPKQSRCTVTGYVEASKVLERVKSTGKAAEMW**PYVPYTMTTYPYVGGAYDKKAPAGFVRGNPAAMADPSAPEVRYMTMFSDENVDSCSIM* |

[1] https://rice.uga.edu/
[2] https://rapdb.dna.affrc.go.jp/
[3] HMA domain is in **bold**. MxCxxC motif (in the HMA domain) is **bold underlined**. CααX motif is underlined.
[4] RAP-DB annotation matches the MSU .1 annotation. The MSU .2 annotation is supported by RNAseq data. No RAP-DB annotation matches MSU.2.
[5] RAP-DB annotation is missing the start codon and first exon.

**Supplementary Table S2. Rice proteins interacting with AVR-Pia in the Y2H screen**

Results of a Y2H screen using BD:AVR-Pia as bait and a cDNA library from the rice cultivar CO39. The number of positive clones with insert sequences (cloned in frame with Gal4 AD) corresponding to the designated proteins are shown. * Potential false positives found in >50% of the Y2H screens performed in the lab using various bait proteins.

| Identified cDNA (MSU/RGAP ID) | Encoded rice protein | Number of clones | Comment |
|---|---|---|---|
| LOC_Os04g48310.1 | RING-H2 finger protein, putative, expressed | 117 | * |
| LOC_Os01g60350.1 | expressed protein | 11 | * |
| LOC_Os02g22130.1 | expressed protein | 8 | * |
| LOC_Os01g13760.1 | dnaJ domain containing protein, expressed | 3 | |
| LOC_Os01g48280.1 | ubiquitin-conjugating enzyme, putative, expressed | 3 | |
| LOC_Os03g16860.2 | DnaK family protein, putative, expressed | 3 | |
| LOC_Os04g37619.1 | zeaxanthin epoxidase, chloroplast precursor, putative, expressed | 3 | |
| LOC_Os05g50710.1 | late embryogenesis abundant protein, putative, expressed | 3 | * |
| LOC_Os07g40580.1 | eukaryotic translation initiation factor 5A, putative, expressed | 3 | |
| LOC_Os12g14070.1 | DnaK family protein, putative, expressed | 3 | |
| LOC_Os01g05060.1 | mitochondrial glycoprotein, putative, expressed | 2 | |
| LOC_Os01g06660.1 | thiamine pyrophosphate enzyme, C-terminal TPP binding domain containing protein, expressed | 2 | |
| LOC_Os02g52290.1 | peptidyl-prolyl cis-trans isomerase, FKBP-type, putative, expressed | 2 | * |
| LOC_Os03g16860.1 | DnaK family protein, putative, expressed | 2 | |
| LOC_Os05g03630.1 | dnaJ domain containing protein, expressed | 2 | |
| LOC_Os05g38370.1 | peptidyl-prolyl cis-trans isomerase, FKBP-type, putative, expressed | 2 | |
| LOC_Os06g04290.1 | S10/S20 domain containing ribosomal protein, putative, expressed | 2 | |
| LOC_Os09g24210.1 | expressed protein | 2 | |
| LOC_Os01g13470.1 | KH domain containing protein, putative, expressed | 1 | |
| LOC_Os01g62610.1 | peptidyl-prolyl cis-trans isomerase, FKBP-type, putative, expressed | 1 | |
| LOC_Os02g02410.1 | DnaK family protein, putative, expressed | 1 | |
| LOC_Os02g08380.1 | CR084 protein, putative, expressed | 1 | |
| LOC_Os02g49150.3 | RNA polymerase Rpb4, putative, expressed | 1 | |
| LOC_Os02g57305.1 | disease resistance protein, putative, expressed | 1 | |
| LOC_Os03g38640.1 | expressed protein | 1 | |
| LOC_Os03g61630.2 | WD domain, G-beta repeat domain containing protein, expressed | 1 | |
| LOC_Os04g01780.1 | uncharacterized ACR, COG1399 family protein, expressed | 1 | |
| LOC_Os04g39560.3 | expressed protein | 1 | |
| LOC_Os05g23740.1 | DnaK family protein, putative, expressed | 1 | |
| LOC_Os07g32380.1 | protein phosphatase 2C, putative, expressed | 1 | |
| LOC_Os07g41810.4 | stress responsive A/B Barrel domain containing protein, expressed | 1 | |
| LOC_Os09g09830.1 | heavy-metal-associated domain-containing protein, putative, expressed | 1 | OsHIPP21 |
| LOC_Os09g36770.2 | NTMC2Type1.2 protein, putative, expressed | 1 | |
| LOC_Os10g08930.1 | S10/S20 domain containing ribosomal protein, putative, expressed | 1 | |
| LOC_Os11g47760.4 | DnaK family protein, putative, expressed | 1 | |
| LOC_Os12g38170.1 | osmotin, putative, expressed | 1 | |

**Supplementary Table S3. Thermodynamic parameters obtained from ITC experiments.**

| | | $K_D$ (M) | ΔH (kcal/mol) | ΔG (kcal/mol) | -TΔS (kcal/mol) | N (sites) |
|---|---|---|---|---|---|---|
| **OsHPP09-HMA & AVR-Pia** | **1** | $1.15 \times 10^{-7} \pm 1.42 \times 10^{-8}$ | -13.7 ± 0.175 | -9.47 | 4.27 | 0.972 ± 0.0052 |
| | **2** | $1.25 \times 10^{-7} \pm 3.02 \times 10^{-8}$ | -11.2 ± 0.277 | -9.42 | 1.79 | 0.859 ± 0.0098 |
| | **3** | $1.85 \times 10^{-7} \pm 5.09 \times 10^{-8}$ | -13.3 ± 0.436 | -9.19 | 4.08 | 0.528 ± 0.01 |
| **OsHPP10-HMA & AVR-Pia** | **1** | $5.26 \times 10^{-6} \pm 1.47 \times 10^{-6}$ | -3.46 ± 0.292 | -7.2 | -3.74 | 1.18 ± 0.055 |
| | **2** | $6.51 \times 10^{-6} \pm 2.04 \times 10^{-6}$ | -3.47 ± 0.345 | -7.08 | -3.61 | 1.17 ± 0.066 |
| | **3** | $7.47 \times 10^{-6} \pm 3.69 \times 10^{-6}$ | -3.18 ± 0.644 | -7 | -3.81 | 1.4 ± 0.177 |
| **OsHPP11-HMA & AVR-Pia** | **1** | $7.72 \times 10^{-7} \pm 2.33 \times 10^{-7}$ | -5.7 ± 0.263 | -8.34 | -2.64 | 0.697 ± 0.018 |
| | **2** | $9.25 \times 10^{-7} \pm 1.78 \times 10^{-7}$ | -5.63 ± 0.182 | -8.23 | -2.6 | 0.727 ± 0.012 |
| | **3** | $9.29 \times 10^{-7} \pm 1.83 \times 10^{-7}$ | -6.28 ± 0.192 | -8.23 | -1.95 | 0.749 ± 0.013 |
| **OsHIPP21-HMA & AVR-Pia** | **1** | $1.29 \times 10^{-6} \pm 1.70 \times 10^{-7}$ | -5.66 ± 0.145 | -8.03 | -2.37 | 0.576 ± 0.0083 |
| | **2** | $1.97 \times 10^{-6} \pm 4.81 \times 10^{-7}$ | -4.84 ± 0.3 | -7.79 | -2.95 | 0.533 ± 0.017 |
| | **3** | $2.33 \times 10^{-6} \pm 3.83 \times 10^{-7}$ | -7.04 ± 0.271 | -7.68 | -0.642 | 0.789 ± 0.016 |

**Supplementary Table S4. X-ray data collection and refinement statistics for OsHPP09-HMA / AVR-Pia**

| | **OsHPP09-HMA / AVR-Pia** |
|---|---|
| **Data collection statistics** | |
| Wavelength (Å) | 0.9655 |
| Space group | $P\,6_1\,2\,2$ |
| Cell dimensions: | |
| *a*, *b*, *c* (Å) | 93.67, 93.67, 72.87 |
| α, β, γ (°) | 90.00, 90.00, 120.00 |
| Resolution (Å)* | 46.83-1.65 (1.68-1.65) |
| $R_{meas}$ (%)# | 4.5 (105.6) |
| $R_{\mathrm{merge}}$ (%)# | 4.0 (96.2) |
| $I/\sigma I$# | 19.2 (1.9) |
| Completeness (%)# | 99.8 (100.0) |
| Unique reflections# | 23168 (1125) |
| Redundancy# | 5.6 (5.9) |
| $CC^{(1/2)}$ (%)# | 99.9 (83.7) |
| **Refinement statistics** | |
| Resolution (Å) | 46.88-1.65 (1.69-1.65) |
| $R_{\mathrm{work}}/R_{\mathrm{free}}$ (%)^ | 17.6/20.2 (29.1/30.4) |
| No. atoms (Protein) | 2125 |
| No. atoms (Ligand/ion) | 10 |
| No. atoms (Water) | 194 |
| B-factors (Protein) | 35.8 |
| B-factors (Ligand/ion) | 33.5 |
| B-factors (Water) | 43.7 |
| R.m.s. deviations:^ | |
| Bond lengths (Å) | 0.0145 |
| Bond angles (°) | 2.243 |
| Ramachandran plot (%): ** | |
| Favoured | 99.2 |
| Allowed | 0.8 |
| Outliers | 0.0 |
| Rotamer outliers (%) ** | 2.65 |
| Clashscore ** | 1.39 |
| MolProbity Score ** | 1.19 |

*The highest resolution shell is shown in parenthesis.

#As calculated by Aimless, ^As calculated by Refmac5, **As calculated by MolProbity

**Supplementary Table S5. Comparison of the binding interfaces of complexes between HMA domains and MAX effectors.**

Interface analysis was performed using qtPISA[4].

| | | | | | | |
|---|---|---|---|---|---|---|
| Complex | **HMA domain** | OsHPP09 | OsHIPP19 | OsHIPP43 | Pikp-1 | RGA5 |
| | **MAX effector** | AVR-Pia | AVR-PikF | HMA PWL2 | AVR-Pia | AVR1-CO39 |
| | **PDB accession code** | 9RSV | 7B1I | 8R7A | 6Q76 | 5ZNG |
| Interface parameter | **Interface area (Å²)** | 521.6 | 1044.3 | 1965.2 | 460.7 | 492.8 |
| | **Solvation energy (kcal/mol)** | -1.9 | -0.1 | -7.6 | -4.7 | -4.6 |
| | **Binding energy (kcal/mol)** | -7.5 | -11.3 | -27.8 | -8.6 | -7.3 |
| | **Hydrophobic P-value** | 0.5066 | 0.6998 | 0.3771 | 0.3701 | 0.3317 |
| | **Hydrogen bonds** | 10 | 17 | 38 | 7 | 6 |
| | **Salt bridges** | 3 | 10 | 9 | 2 | 0 |
| | **Disulphide bonds** | 0 | 0 | 0 | 0 | 0 |
